# CyclicCAE: A Conformational Autoencoder for Efficient Heterochiral Macrocyclic Conformational Sampling

**DOI:** 10.1101/2025.02.21.639569

**Authors:** Andrew Powers, P. Douglas Renfrew, Parisa Hosseinzadeh, Vikram Khipple Mulligan

## Abstract

Peptide macrocycles are a promising therapeutic class. The inclusion of heterochiral and non-natural amino acids allows far more folds and functions to be accessed, but creates challenges for rational design – particularly for sampling plausible mainchain conformations. We developed a conformational autoencoder called CyclicCAE to rapidly generate energetically favourable macrocycle scaffolds for heterochiral design and structure prediction. Given the absence of large, available macrocycle datasets, we created a custom dataset *in silico* using physics-based simulation methods. Trained on this, CyclicCAE produces energetically stable mainchain conformations and designable scaffolds more rapidly than the current state-of-the-art method, the Rosetta software suite’s Generalized Kinematic Closure (GeneralizedKIC) method. We show that, despite being trained exclusively on synthetic data, CyclicCAE accurately captures the conformations accessible to peptide macrocycles found in the Protein Data Bank, with considerable speed advantages over GeneralizedKIC. We also demonstrate that CyclicCAE enables users to perform energy minimization in isolation or in a target-bound context, to generate structurally similar or diverse outputs by Markov Chain Monte Carlo sampling, and to conduct inpainting with fixed motifs. This method, which we release under a free and open source licence, will accelerate macrocycle design pipelines, speeding the development of peptide therapeutics.

**Conventions:** In this work, vectors are represented with an arrow (*e*.*g*. 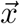), and matrices or higher-order tensors with a bar (*e.g*. 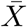). Entries in vectors are represented with a subscript index (*e.g. c*_*i*_). A hat indicates an expected or desired value (*e.g*. 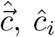). The Greek letters *θ* and *ϕ* are used to represent bond angles and dihedral angles, respectively, while *c* is used for a generic degree of freedom.

## Introduction

A promising class of therapeutics, peptide macrocycles present larger surfaces for specific target recognition than do small molecules, while retaining greater potential for permeability, immune system evasion, and shelf stability than large proteins [1, 2]. Cyclization can enhance conformational stability, provide greater protection against proteolytic degradation, and improve membrane permeability [3, 4, 5, 6, 7, 8, 9, 10]. Solid-phase peptide synthesis enables the incorporation of not only the 20 canonical amino acids, but of non-canonical amino acids (NCAAs), which can increase resistance to proteolysis, provide structural rigidity in more diverse conformations, or confer other desired properties, increasing the chances of finding new, functional peptide folds for a given application [5, 11, 12, 13].

Machine learning (ML) is a broad term for computational approaches in which performance on a task, assessed by some metric, improves on exposure to data. Recently, deep learning (DL), the subdiscipline of ML based around deep neural networks, has revolutionized protein structure prediction and enhanced canonical protein design [14, 15, 16, 17]. DL has benefitted NCAA peptide macrocycle design and validation less, however. Current approaches that adapt existing DL models for macrocycles, such as AfCycDesign [18], RFPeptide [19], and HighFold [20] have shown success predominantly for l-homochiral macrocycles built from the same canonical *α*-amino acid building-blocks as the Protein Data Bank (PDB) proteins used in the training set.

These models poorly generate or predict structures of heterochiral macrocycles, for which large databases of experimental training data are not available. The StrEAMM and StrEAMM-thioether models [21, 22], which were trained on molecular dynamics (MD) simulations of heterochiral penta- and hexapeptides, are the exception. However, due to the expense of the MD simulations generating the training data, these models do not currently extend beyond six-residue macrocycles.

Physics-based methods that perform semi-random sampling in a manner that is less biased by prior knowledge, such as the peptide macrocycle workflows implemented in the Rosetta software suite, are currently the state of the art for designing and predicting structures of heterochiral and NCAA macrocycles (reviewed in [23, 24, 25]). Since these methods sample without simulating time-evolution between states, they produce conformational ensembles at a fraction of the computational cost of MD simulations. These methods have been successfully applied to create a wide range of well-folded macrocycles [26, 11, 27, 28, 29], including macrocycles that are able to bind to and inhibit targets of therapeutic relevance, such as the New Delhi metallo-*β*-lactamase (NDM1) [12] or histone deacetylases [13]. The current method for macrocycle mainchain conformational sampling in Rosetta is the Generalized Kinematic Closure (`GeneralizedKIC`) algorithm [26, 30] (see Supplementary Information section S1.2), which solves analytically for a subset of the internal degrees of freedom of a chain of atoms in order to connect the start and end of a loop or a closed cycle [31, 32, 26]. Generating a full conformational ensemble for an 8-residue macrocycle can require sampling up to 50,000 conformations [11]. Unfortunately, this stochastic method is prone to spending time sampling energetically unfavourable regions of the conformational landscape. Repeatedly carrying out near-exhaustive landscape exploration to find scaffolds for a large batch of designs, then repeating the sampling again to validate each designed sequence, is costly: a faster means of sampling from the plausible set of conformations is needed.

Rather than using DL models to learn patterns from large databases of experimental data, we sought to use them as fast function approximators, to learn to rapidly (albeit approximately) perform a task is costlier to perform by physics-based simulation. Here, we present a new DL model called CyclicCAE (short for the *Cyclic C* onformational *A*uto*E* ncoder). Trained on an ensemble of peptide macrocycle conformational samples using `GeneralizedKIC`, CyclicCAE permits new conformations to be cheaply sampled from the distribution of accessible, low-energy closed conformations. The autoencoder architecture is based on one-dimensional convolutions, which permits inference at far lower computational expense than would be required for a diffusion model. This was chosen to maximize the speedup while still ensuring that the distribution of accessible conformations could be accurately learned. After training once on conformational samples produced at greater expense, a trained CyclicCAE instance may be used many times, for many drug design problems, yielding considerable reduction in the computational cost of macrocycle drug design. CyclicCAE also offers versatility advantages over `GeneralizedKIC`, permitting inpainting against desired motifs. The loss function used for training ensures that conformations with low root mean squared deviations (RMSDs) are mapped to points nearby in the latent space, facilitating local conformational searches to diversify from a promising starting point. Since the decoder limb is also a regression model that predicts conformational energy in addition to generating a conformation, it is possible to descend the energy gradient in the latent space, effectively relaxing mainchain conformations on the manifold of accessible conformations, permitting more extensive relaxation while preserving favourable features like hydrogen bond networks. A binding pocket context term added to the loss function used at inference time makes CyclicCAE highly suitable for macrocycle drug design in binding pockets on target proteins. Given these features, we anticipate that CyclicCAE will be broadly useful for reducing the computational costs of producing initial candidate designs and of validating designs by conformational sampling, thus accelerating macrocycle drug design pipelines.

## Methods

### Model and dataset construction

We implemented a convolutional autoencoder, CyclicCAE, that maps a vector of mainchain degrees of freedom (DoFs) for an *α*-amino acid macrocycle onto a low-dimensional latent space, then converts the latent space representation back into a DoF vector plus predicted conformational energies (**Fig. 1**, with details in **Ext. Data Fig. 1** and in Supplementary Information section S3.1).

**Fig. 1.**
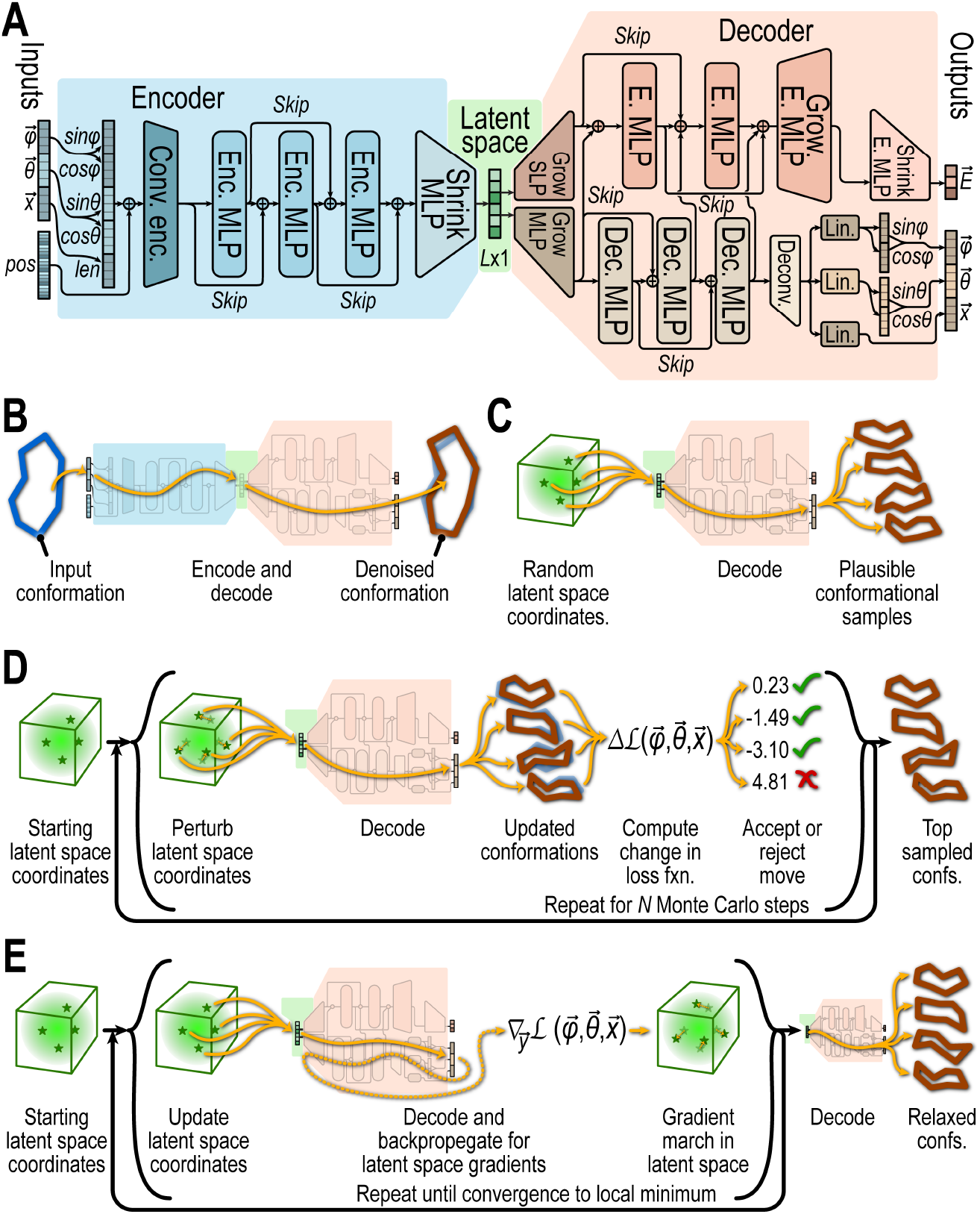
CyclicCAE architecture and usage. (**A**) Architecture of the CyclicCAE deep neural network. CyclicCAE’s encoder limb (blue) accepts vectors of mainchain dihedral angles 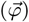, bond angles 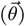, and bond lengths 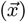, combines these with positional encodings (*pos*), and uses a series of 1D convolutional blocks and multilayer perceptron (MLP) blocks to map these to an *L*-dimensional latent space (green) in a manner that ensures that similar conformations are nearby. It then decodes this in two limbs, one of which uses a series of MLP blocks (pink) to output vectors of energies 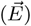, and the other of which uses MLP blocks followed by a deconvolutional block (beige) to reconstruct 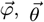, and 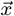 degree of freedom vectors. See **Ext. Data Fig. 1** for details. (**B**) Encoding and decoding to “de-noise” structures, mapping input conformations to the nearest stable mainchain conformation that the model can represent. (**C**) Sampling directly from the latent space and decoding to rapidly generate plausible macrocycle conformations. (**D**) Monte Carlo exploration of the latent space to find plausible conformations that also have a desired property, given a loss function 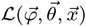 that scores the property as a function of conformation. This loss may include components of 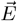 generated by the model. (**E**) Gradient descent minimization in the latent space to optimize a desired property while preserving conformational plausibility. A differentiable loss function 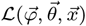 is needed. The gradient 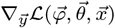 with respect to a latent-space coordinate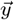 is computed by back-propagation. Since 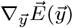 is easily computed, the loss function may include one or more energies predicted by the model.

To train CyclicCAE, we needed a large dataset of peptide macrocycle conformations; however, compared to linear peptides and proteins, there are few experimentally determined heterochiral macrocyclic peptide structures. We therefore generated a synthetic dataset of macrocycle conformations using Rosetta’s `simple_ cycpep_predict` application [11, 26], which uses `GeneralizedKIC` to produce macrocycle conformational ensembles (Supplementary Information section S1.2). We produced 50,000 poly-glycine 6-, 7-, and 8-mer conformations, and 100,000 poly-glycine 9-mer conformations. Glycine is unique among the canonical α-amino acids because it is achiral, and its accessible mainchain dihedral angles are roughly the union of the sets of mainchain conformations accessible to l- and d-amino acids.

The generated structures were then subjected to gradient-descent geometry optimization using the `GAMESS` quantum chemistry package [33, 34] by way of the `RosettaQM` bridge (manuscript in preparation). Final energies were computed with both Rosetta’s `ref2015` force field and with GAMESS using the DFTB semi-empirical quantum chemistry method [35] with SMD implicit organic solvent [36] to promote hydrogen bond formation.

### Assessment of CyclicCAE’s ability to reproduce known peptide conformations

We exhaustively searched the PDB (version current as of 14 January 2026) for experimentally determined structures of peptide macrocycles of 6 to 9 amino acid residues, cyclized by an N-to-C amide bond, free or bound to a target protein. Peptides with multiple cycles (internal crosslinks) were excluded. This yielded 11 examples of 6-mers, 8 examples of 7-mers, 12 examples of 8-mers, and 6 examples of 9-mers. PDB IDs are listed in Supplementary Table S6. Of the peptides found, some were originally computationally designed using `GeneralizedKIC` with an anchor extension approach [12, 13]. In these cases, we used these anchor residues when sampling conformations with CyclicCAE. We encoded and decoded (**Fig. 1B**) all cyclic permutations of all macrocycles found, and we measured the minimum mainchain heavy-atom RMSD of the decoded structure to the input.

### Analysis of efficiency of CyclicCAE for designing stable folds

We sampled 5,000 macrocycle scaffolds using each of `GeneralizedKIC` and CyclicCAE, each running on a single CPU core (AMD Genoa). We also generated 5,000 scaffolds with CyclicCAE on a single GPU (NVIDIA A100). Protocols for producing samples are described in Supplementary Information sections S1.2.1 and S4.5. These scaffolds were then subjected to the same Rosetta `FastDesign` design script, which used the `aa composition` design-centric guidance term [11] to guide the design process to sequence compositions favouring folding, as described in Supplementary Information section S1.5.1. These designs were then filtered to identify good candidates for stability testing. Any peptide that had a non-negative computed energy with the Rosetta `ref2015` energy function [37] was excluded, as was any peptide lacking dor l-proline. Supplementary Table S4 shows the number of designs passing filters. From these, for each scaffold generation method (CyclicCAE or `GeneralizedKIC`) and macrocycle size, a random subset of designs was analyzed with Rosetta’s `simple_cycpep_predict` application to compute the *P*_near_ fold propensity metric, by a protocol described in section S1.3.3. The value of *P*_near_ ranges from 0, for sequences with no propensity to adopt the desired fold, to 1, for sequences that strongly favour the desired fold [11, 26]. Since `simple_cycpep_predict` analysis is expensive and grows more so with macrocycle size, the number of designs analyzed in this way was chosen based on available CPU time. Counts ranged from 465 (9-residue `GeneralizedKIC` designs) to 1,000 (6-residue `GeneralizedKIC` designs), and are shown in round brackets in **Fig. 3A**. Given a desired fold propensity 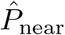, for each scaffold generation method we calculated the expected time 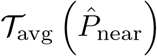 needed to produce at least one design with this fold propensity using Eq. (1) (assuming that the generated scaffolds were subjected to the same `FastDesign` protocol to yield final designs).

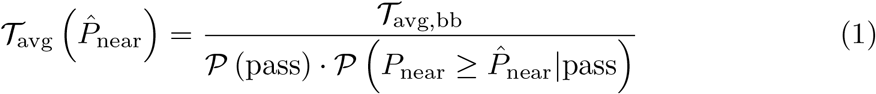

In Eq. (1), *T*_avg,bb_ is the mean time to generate one scaffold by the method, *P* (pass) is the probability that a scaffold generated by the method passes filters after design, and 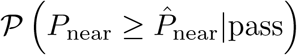 pass is the probability that a scaffold generated by the method that passes filters after design has a *P*_near_ value greater than or equal to the desired value 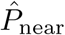.

### Motif inpainting

We developed an approach for using CyclicCAE to generate scaffolds in which 1 to 4 residues are in a desired conformation (such as one taken from a known structure) and the conformations of the rest are chosen to yield a closed macrocycle, with favourable geometry presenting this motif. In analogy to generative AI that fills in missing areas of images, we call this “inpainting”. Our approach involves Monte Carlo searches of the latent space to find conformations with segments approximating the desired motif, followed by modified gradient descent minimization of predicted energy within the latent space with an additional loss term to bring the conformation of the motif-matching residues closer to that of the desired motif. Please see Supplementary Information sections S3.3.2 and S4.7 for full details.

### Sampling macrocycle backbone conformations in the context of a target binding pocket

To design macrocycle drugs, on must sample plausible backbone conformations in the context of a binding pocket on a target protein, seeking backbones that could be stabilized by suitable subsequent choice of sequence. In order to enable macrocycle design for a particular interface or pocket, we added a Lennard-Jones style repulsion term to the CyclicCAE loss function used at inference time to penalize clashes between sampled scaffold atoms and target mainchain or C_*β*_ heavy-atoms. This term is explained in Supplementary Information section S3.3.3. The overall loss function, described in Supplementary Information section S3.3, includes terms penalizing poor terminal bond geometry, high predicted energy, and distortion of the desired binding motif in addition to this pocket context repulsion term. This permits both Monte Carlo searches (Supplementary Information section S4.6) and gradient-descent minimization (Supplementary Information sections S3.4 and S4.4) in the CyclicCAE latent space, yielding macrocycle conformations that are favourable in isolation and which also provide good shape complementarity to a target binding pocket.

We tested this by sampling backbone conformations compatible with binding to the New Delhi metallo-*β*-lactamase 1 (NDM1, PDB ID 6XBF [12]) and to the murine C5a anaphylatoxin chemotactic receptor 1 (C5AR1), a G-protein coupled receptor (GPCR, PDB ID 8HQC [38]). In the case of NDM1 we emulated a previous approach [12], starting with a d-cysteine, l-proline two-residue stub that resembles l-captopril, a small-molecule inhibitor of NDM1. In the case of the GPCR, our starting stub was residue Leu72 of the C5a anaphylatoxin, which in the 8HQC cryo-electron microscopy structure is bound in a tight pocket on the GPCR [38]. We sampled backbones in the binding pocket context using Monte Carlo searches followed by gradient-descent relaxation. During the search and the relaxation steps, we used motif constraints (Supplementary Information section S4.7) to ensure that sampled scaffolds were compatible with each stub, while including the Lennard-Jones term during sampling to find backbone conformations that minimally clashed with the target. We assessed the fraction of backbones that clashed with the target with and without the Lennard-Jones term.

## Results

### CyclicCAE’s latent space encodes macrocycle conformations observed in experimentally determined structures

To be useful, CyclicCAE must first be able to accurately represent real peptide macrocycle conformations. To test the trained model’s accuracy, we passed experimentally-determined macrocycle structures taken from the Protein Data Bank (PDB) through the model (**Fig. 1B**), assessing the reproduction of mainchain conformations matching the inputs. Our test set consisted of a total of 37 x-ray crystal or NMR structures of macrocycles ranging from 6 to 9 residues, none of which were in the simulation-based training data.

Comparing input and output structures from this process, we found median mainchain heavy-atom RMSD values less than 1 Å for 6, 7, and 8-residue macrocycles (**Fig. 2**). For 9-residue macrocycles, the median RMSD increased to 1.8 Å. Among 6-mers, the largest RMSD outlier (1.360 Å) was NMR structure 1SKL (**Fig. 2**-2). This peptide has a sequence with many bulky, hydrophobic NCAA sidechains. Hydrophobic contacts between these stabilize 1SKL in a strained mainchain conformation that CyclicCAE (which has no knowledge of sequence) fails to recognize as low-energy. More than half of the surface of 7-mer RMSD outlier 7BTI (**Fig. 2**-4; 1.577 Å) interacts with a target protein in the x-ray crystal structure, stabilizing it in an unusually open conformation with no internal hydrogen bonds. 8-mer RMSD outlier 6AXI (**Fig. 2**-6; 1.848 Å), which is also a solution NMR structure, has a large flexible region outside of its two contiguous proline residues, which adopts transiently stable conformations.

**Fig. 2.**
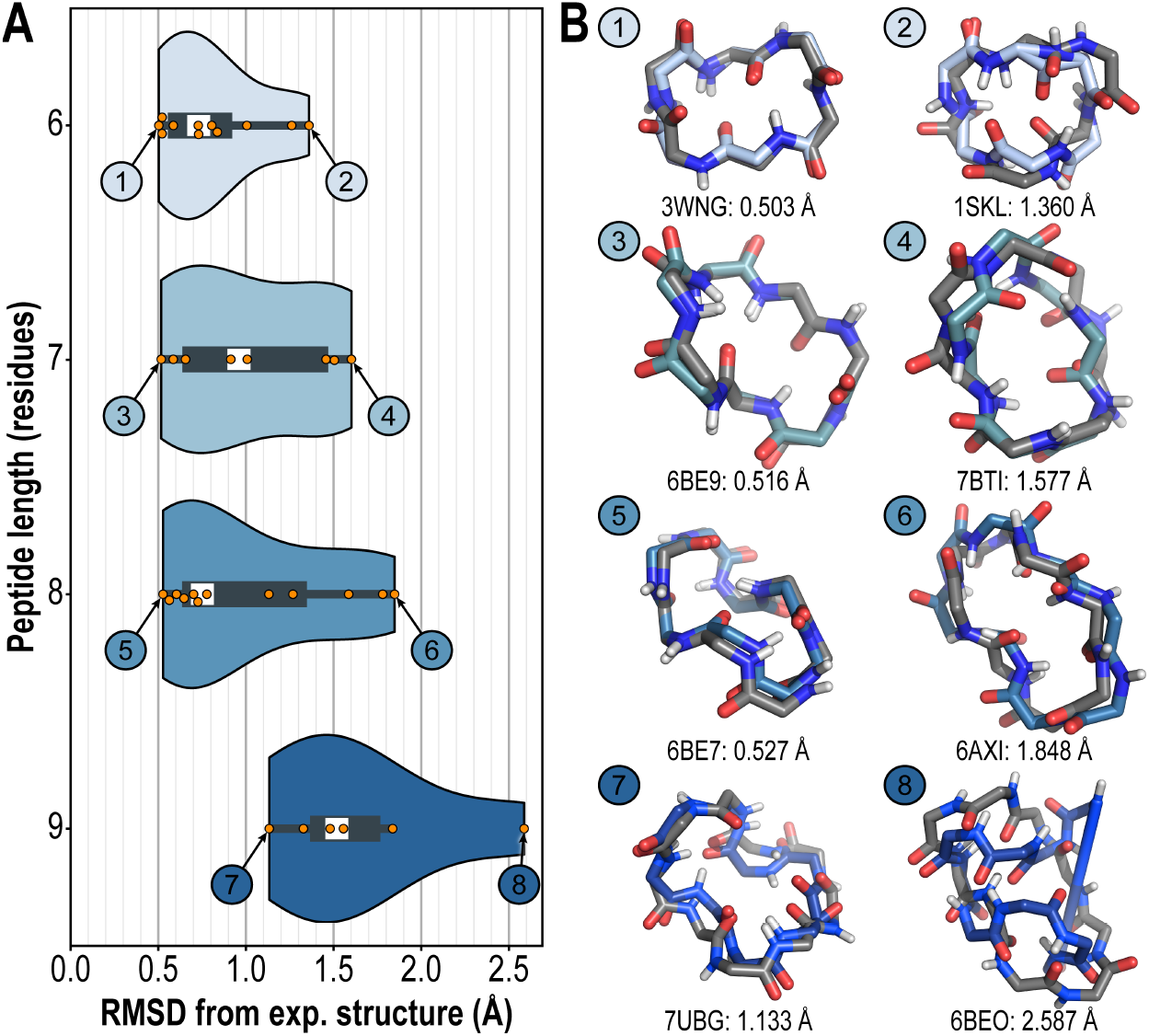
Reproduction of experimentally-determined peptide macrocycle mainchain conformations by CyclicCAE. PDB structures were encoded and decoded. (**A**) Violin plots showing distribution of RMSD values observed, with mean marked in white, for each peptide size. (**B**) Minimum and maximum RMSD peptides for each size, with model output (blues) overlain on the PDB structure (grey). Mainchain heavy-atom RMSD values are shown below each structure pair.

Interestingly, 9-mer RMSD outlier 6BEO (**Fig. 2-8**; 2.587 Å) does appear to have a stable structure in solution based on solution NMR analysis [11], yet the encoder maps this to a region of the latent space which, when decoded, results in extremely distorted terminal bond geometry. Thus in all but one of these cases in which CyclicCAE most poorly encodes the conformation, either intrinsic disorder, or stabilization by sequence features or the peptide environment, are likely. To understand the exceptional 9-mer case, we next examined whether minimization of the CyclicCAE decoder’s estimate of the energy by gradient descent in the latent space improved the decoded structures (**Fig. 1D**; Supplementary Information section S4.4). To enable latent-space gradient descent without distortion of the cyclic geometry, we implemented a loss function based on the energy estimate produced by the model plus a penalty for distorted terminal bond geometry (Supplementary Information section S3.2.2 and **Ext. Data Fig. 2**). As shown in **Ext. Data Fig. 3**, 10 steps of gradient descent minimization had little effect on the mean or variance of the distribution for 6, 7, or 8-mer peptides, only producing a large reduction in the mean RMSD (from 1.5 to 1.3 Å) and the largest outlier (from 2.587 to 1.935 Å) for 9-mer peptides. Indeed, in this case, structure 6BEO changes from being the worst outlier to the *best* peptide, with an RMSD of only 1.221 Å. This indicates that the latent space and decoder limb robustly represent this conformation, and that the problem is in the encoder limb.

The new highest-RMSD 9-mer peptide with gradient-descent refinement is 6WI3 (**Ext. Data Fig. 3-8**; 1.935 Å RMSD with latent-space gradient descent refinement). The x-ray crystal structure features three copies of this peptide bound to a target protein. One of the copies was unresolved, while the other two differ by 1.907 Å mainchain heavy-atom RMSD. Both resolved conformations show no internal intra-mainchain hydrogen bonds. Combined with the conformational heterogeneity, this suggests that this peptide has no unique stable fold in isolation.

### CyclicCAE accelerates the design of stable peptide macrocycle folds

We next tested the performance of the decoder portion of CyclicCAE in a peptide design pipeline, sampling from the latent space to generate initial mainchain conformations passed to Rosetta’s `FastDesign` protocol [26], and comparing this to the same protocol using `GeneralizedKIC` to produce initial mainchain samples. `GeneralizedKIC` samples possible conformations of a macrocycle mainchain biased by the Ramachandran preferences of each amino acid residue, but cannot draw directly from the distribution of low-energy states for the whole macrocycle [11, 26]. Since CyclicCAE was trained to predict overall energy, with gradient-based refinement, it is expected to have advantages over `GeneralizedKIC` in both speed and likelihood of sampling productive conformations. We generated 5,000 macrocycle conformations using each of `GeneralizedKIC` (Supplementary Information section S1.2) and CyclicCAE (**Fig. 1C,D**; Supplementary Information section S4.5), then passed them all through the same `RosettaScripts` design script (Supplementary Information section S1.5). After filtering, we analysed a subset of designs produced by each method, using Rosetta’s `simple_cycpep_predict` to compute the *P*_near_ fold propensity metric (see Supplementary Information section S1.3.3 and [26]).

We found that for 7-, 8-, or 9-residue macrocycles, `FastDesign`-based sequence design was much more likely to produce designs with high *P*_near_ values ≥ (0.8) if performed on CyclicCAE-generated scaffolds instead of `GeneralizedKIC`-generated scaffolds (**Fig. 3A**). This resulted in a one to two order of magnitude reduction in the sampling time needed on a single CPU core to produce a high-quality design with *P*_near_ ≥ 0.8 (**Fig. 3B**). For 6-mers, likelihood of high *P*_near_ after sequence design was comparable for scaffolds produced by either method. The time to produce *N* samples was 2.2*×* (6-mers), 2.9*×* (7-mers), 3.3*×* (8-mers), or 3.5*×* (9-mers) less with CyclicCAE than with `GeneralizedKIC` on a single CPU core. Since CyclicCAE is a neural network, GPU execution yields sampling times 5.7*×* (6-mers), 7.3*×* (7-mers), 8.2 *×* (8-mers), or 7.1*×* (9-mers) faster than `GeneralizedKIC` sampling times on a single CPU core (insets in **Fig. 3B**).

**Fig. 3.**
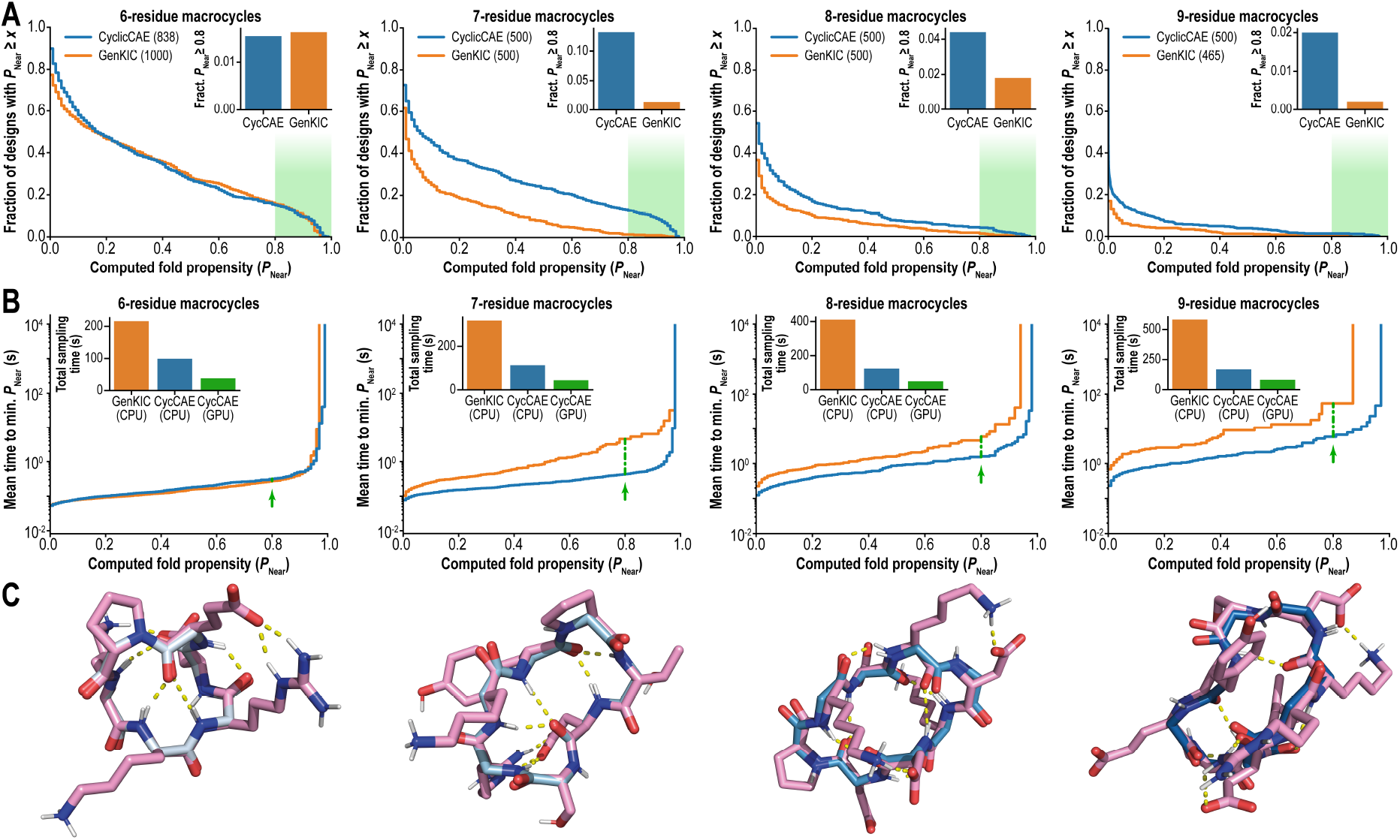
CyclicCAE performance in a peptide design pipeline. (**A**) Cumulative distribution of computed peptide fold propensity (*P*_near_, from conformational sampling using Rosetta’s `simple cycpep predict` application) for peptide macrocycles designed with Rosetta’s `FastDesign` protocol using scaffolds either sampled with Rosetta’s `GeneralizedKIC` algorithm (orange) or with CyclicCAE (blue). Macrocycles with *P*_near_ ≥ 0.8 are considered to fold stably (green shading). For peptides larger than 6 residues, CyclicCAE produces a considerably larger fraction of samples which can be stabilized with a suitable choice of sequence (insets). (**B**) Mean sampling walltime on a single CPU core needed to produce at least one sampled conformation that yields *P*_near_ greater to or equal than a desired value after `FastDesign`-based sequence design. Insets show total time needed to sample 5,000 mainchain conformations with GenKIC on a single AMD Genoa CPU core (orange) or with CyclicCAE on a single AMD Genoa CPU core (blue) or on an NVIDIA A100 GPU (green). To achieve desired *P*_near_ ≥ 0.8, CyclicCAE on a single CPU core provides comparable performance to `GeneralizedKIC` on a single CPU core for 6-mers, and speedups of one to two orders of magnitude for 7-mers, 8-mers, and 9-mers (green arrows and dashed green lines). (**C**) Representative designed structures with *P*_near_ ≥ 0.8, using scaffolds generated by CyclicCAE. White to cyan: initial scaffold from CyclicCAE. Pink: final relaxed structure following sequence design (`FastDesign`).

**Fig. 3C** shows representative high *P*_near_ final designs produced with CyclicCAE-generated scaffolds. The original mainchain conformations produced by CyclicCAE are shown overlain on the relaxed final designs. The rounds of relaxation performed by `FastDesign` result in very little change in the mainchain conformation, indicating that CyclicCAE’s initial output was an energetically favourable conformation.

### Generating macrocycles from linear motifs with conformational motif inpainting

Conformational motif inpainting [39, 40] involves generating plausible conformations of residues adjacent to a region of a macromolecule that is in a desired, fixed conformation. In the context of peptide macrocycle design, this allows known linear functional or binding motifs to be stabilized in a minimal macrocyclic scaffold. Traditional conformational motif inpainting generates Cartesian atom coordinates, whereas our DoF-based motif inpainting generates mainchain torsion angles, bond angles, and bond lengths. Our CyclicCAE-based approach is described in Supplementary Information section S4.7.

Using x-ray crystal structures of a protein loop from the N-formyl peptide receptor 2 (FPR2, PDB ID 6LW5), the linear glucagon-like peptide-1 (GLP1, PDB ID 6×18), and a mixed-chirality peptide macrocycle inhibitor of NDM1 (NDM1i-1G, PDB ID 6XBF) as source scaffolds, we extracted contiguous motifs ranging from 1 to 4 residues, and used these to generate 6-to 9-residue macrocycles presenting these motifs. We generated 6-mers presenting 1 to 2 residue motifs, 7- and 8-mers presenting 1 to 3 residue motifs, and 9-mers presenting 1 to 4 residue motifs. As shown in **Fig. 4**, CyclicCAE is robustly able to generate macrocycles presenting 1 to 2 residue motifs, with motif mainchain heavy-atom RMSDs of 0.2 Å or less. With the exception of 9-mers presenting the loop region motif, times to propose 1,000 scaffolds on a single CPU core were less than 100 s. 3-residue motifs showed slightly higher RMSDs (approaching 0.5 Å in some cases), but these still closely matched the desired motif, with compute times for 1,000 proposed scaffolds under 100 s for 7- and 8-mers, and under 200 s for 9-mers. 4-residue motifs, which took longer to compute (showing times over 200 s), showed average motif RMSDs exceeding 1.5 Å in two of three cases. This larger deviation is unsurprising given that one would expect it to be harder to find larger motifs in the embedded accessible conformations of a macrocycle. In the third case, a motif taken from a peptide macrocycle instead of from a linear peptide or protein, CyclicCAE proposed scaffolds with lower motif RMSDs (0.9 Å on average). This is unsurprising since this motif is known *a priori* to be compatible with at least one macrocycle conformation. For smaller contiguous motifs, CyclicCAE proves able to propose macrocycle scaffolds to present desired motifs robustly and rapidly.

**Fig. 4.**
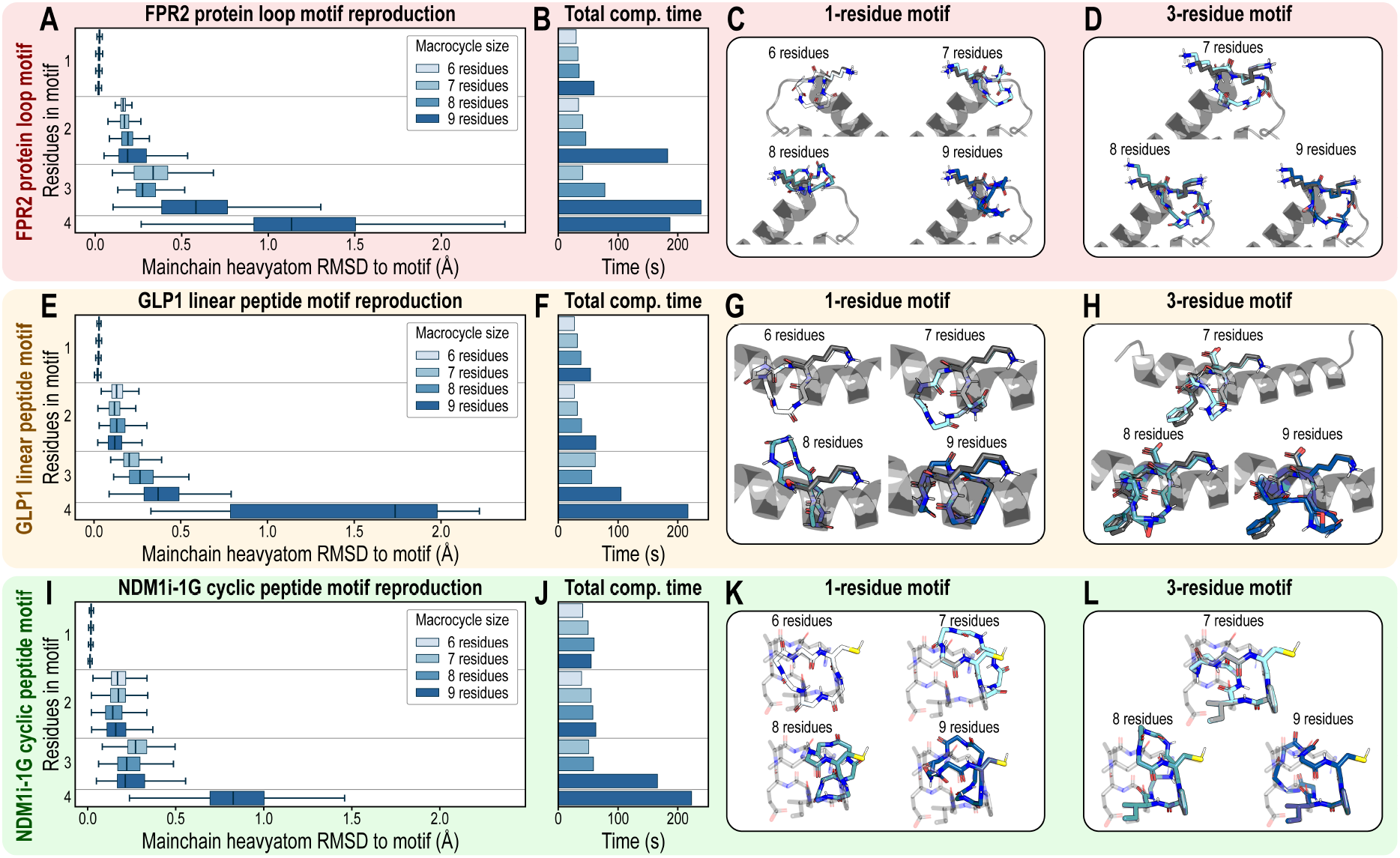
Use of CyclicCAE to sample conformations compatible with an input motif by inpainting. Test motifs were taken from a protein loop in the N-formyl peptide receptor 2 (FPR2, pink, panels **A-D**), from the glucagon-like peptide 1 (GLP1, yellow, panels **E-H**), and from a cyclic peptide inhibitor of NDM1 (NDM1i-1G, green, panels **I-L**). (**A, E, I**) Distribution of mainchain heavy-atom RMSD values between input motif and output peptide macrocycle, for motifs of 1-4 residues and output macrocycles of 6-9 residues. 1,000 outputs were produced for each input. (**B, F, J**) Total computing time on a single AMD Genoa CPU core to generate 1,000 outputs for the inputs shown in the previous column. (**C, G, K**) Representative output structures across a range of output macrocycle sizes (shades of cyan) given a single-residue motif (grey sticks). Scaffolds providing the motifs are shown as grey cartoons. (**D, H, L**) Representative output structures given a three-residue motif. Colours are as in the previous column.

We also confirmed that CyclicCAE can find known scaffold conformations able to present known motifs. We used CyclicCAE to generate scaffolds able to present a one-residue motif taken from each of six experimentally-determined macrocycle structures from the PDB. In each case, CyclicCAE was able to propose scaffolds with mainchain heavy-atom RMSDs of less than 1 Å to the native structure, as well as other, diverse macrocycle conformations that could also present the motif (**Ext. Data Fig. 4**).

### CyclicCAE samples plausibly binding-competent conformations in the context of target binding pockets

Macrocycle drug design pipelines demand means of proposing macrocycle conformations compatible with binding to targets of therapeutic interest. We next tested CyclicCAE’s ability to sample and to relax scaffolds in the context of a target protein’s binding pocket. For this task, we used two targets: NDM1 and the C5AR1 G-protein coupled receptor (GPCR). The latter was used since GPCRs are of high interest as drug targets, with tight pockets that make it particularly challenging for existing methods to generate stable macrocycles that do not clash with pocket sidechain or backbone atoms. We generated four sets of 100 scaffolds, one for each target with or without the Lennard-Jones term described in Supplementary Information section S3.3.3.

When scaffolds were generated and relaxed by CyclicCAE without the Lennard-Jones term penalizing clashes with pocket atoms, 4% of NDM1 peptides and 25% of GPCR peptides had clashes with the target, based on a clash threshold of 0.5 Å between any peptide heavy-atom and target backbone heavy-atom or C_*β*_ atom. When the stringency of this requirement was increased, so that the clash distance threshold was 1.0 Å, then 35% of NDM1 peptides and 61% of GPCR peptides clashed with their target. In contrast, inclusion of the pocket context (and the Lennard-Jones term) during relaxation by CyclicCAE resulted in *none* of the NDM1 peptides and only 6% of the GPCR peptides clashing with their targets by the 0.5 Å criterion, and only 3% of NDM1 peptides and 19% of GPCR peptides clashing by the 1 Å criterion (**Fig. 5A-B**). Generated conformational ensembles were compact and shaped to the binding pocket with the Lennard-Jones context term enabled (**Fig. 5C-H**). This is a clear demonstration of the efficacy of CyclicCAE for rapidly finding macrocyclic backbones that are plausibly compatible with any given target binding pocket, which could serve as inputs to a subsequent `FastDesign` sequence design script as part of a macrocycle drug design pipeline.

**Fig. 5.**
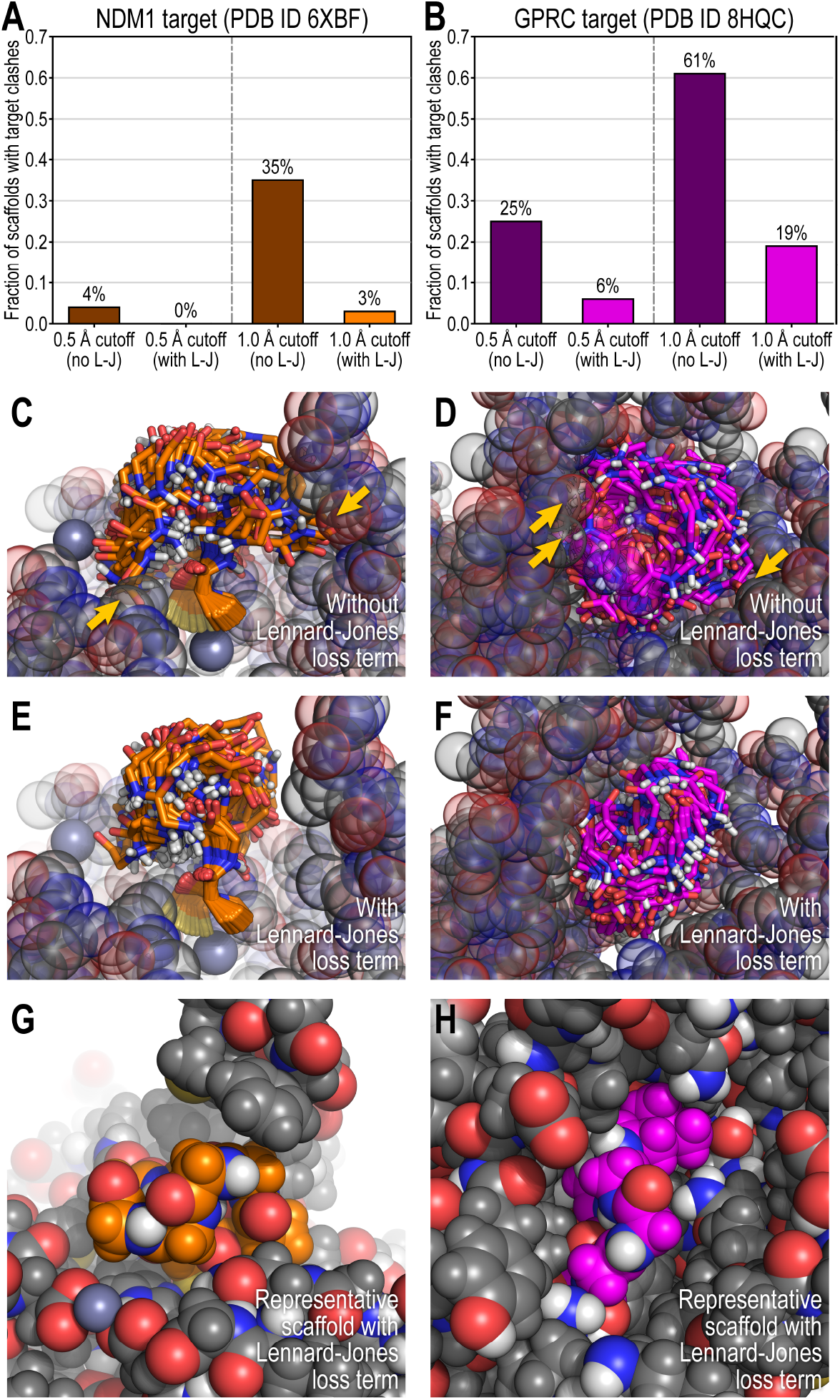
Use of CyclicCAE to generate plausible macrocycle scaffolds in binding pockets. NDM1 (PDB ID 6XBF) and the C5AR1 GPCR (PDB ID 8HQC) were used as targets, and scaffolds were sampled by Monte Carlo sampling and gradient descent in the latent space, minimizing the loss function shown in Eq. (S20) in the Supplementary Information, which contains terminal geometry, energy estimate, motif geometry, and Lennard-Jones (pocket context) terms. (**A-B**) Fraction of generated macrocycle scaffolds that clash with NDM1 (**A**, orange) or GPCR (**B**, magenta) target mainchain or C_*β*_ heavy-atoms without (dark bars) and with (light bars) the Lennard-Jones term in the loss function to penalize clashes. (**C-F**) Generated scaffold ensembles (orange, magenta) for NDM1 (**C, E**) or the GPCR (**D, F**). Target mainchain and C_*β*_ heavy-atoms are shown as grey spheres. Inclusion of the Lennard-Jones term (**E, F**) produes more compact ensembles that better fit target pockets, as compared to generation without this term (**C**,**D**), which yields visible target clashes (yellow arrows). (**G-H**) Representative scaffolds (orange, magenta) produced with the Lennard-Jones term to target NDM1 (**G**) or the GPCR (**H**), showing snug, clash-free fits in their binding pockets. All target atoms are shown (grey).

## Discussion

CyclicCAE, our deep-learning model trained on simulation output for the generation of heterochiral macrocyclic peptide scaffolds, is intended to accelerate peptide macrocycle drug design pipelines. While previous simulation-based approaches such as `GeneralizedKIC` provided general means of exhaustively sampling conformations of macrocycles, repeated simulation-based sampling carried heavy computational cost. By learning the distribution of conformations produced by a single expensive simulation, CyclicCAE provides a fast means of repeatedly sampling from this distribution in macrocycle drug design pipelines. In order to be useful for this task, CyclicCAE must meet three criteria. First, CyclicCAE must be *accurate*, correctly reproducing the plausible low-energy conformations accessible to macrocycles. Second, CyclicCAE must be *fast*, permitting more rapid sampling of useful, productive macrocycle conformations than does `GeneralizedKIC`. And third, CyclicCAE must be *versatile*, and usable in the same design contexts as `GeneralizedKIC` — that is, not only for unconstrained sampling of macrocycle conformations, but also for discovering binding-competent conformations able to present desired interaction motifs to a target, and which are shaped to fit target binding pockets.

To assess accuracy, the first criterion, we showed that CyclicCAE correctly reproduces the plausible low-energy conformations accessible to macrocycles. Despite being trained exclusively on simulated conformational ensembles of poly-glycine, CyclicCAE’s latent space captures experimentally observed conformations of mixed-chirality peptides on which it was not directly trained (**Fig. 2**). CyclicCAE’s estimate of conformational energy can be further used to refine sampled conformations (**Ext. Data Fig. 3**). In the context of a peptide design pipeline, we also found that, for macrocycles larger than 6-mers, CyclicCAE-generated conformations were far more likely to yield high stability after sequence design (*P*_near_ ≥ 0.8) than were `GeneralizedKIC`-generated conformations (**Fig. 3A**, **inset**).

To ensure speed, the second criterion, we chose an autoencoder architecture that allows more rapid sampling than would more computationally costly approaches like diffusion models. We assessed speed by showing that CyclicCAE samples structurally diverse, low-energy macrocycle conformations 2 to 3 times faster than `GeneralizedKIC`, even on a single CPU core, and that GPU parallelism increases the speedup to about an order of magnitude faster than `GeneralizedKIC` (**Fig. 3B**, **inset**). Coupled with the higher probability of CyclicCAE-generated conformations being productive (yielding *P*_near_ ≥ 0.8 after sequence design), this meant that, compared to `GeneralizedKIC`, CyclicCAE yielded an order of magnitude reduction in the sampling time required to find a productive conformation on a single CPU core for 7-mers and 9-mers, and about half an order of magnitude reduction for 8-mers. Since a larger fraction of sampled conformations yield productive designs, this also means a reduction in the average time spent in downstream stages of the design pipeline per successful design generated – particularly in the computationally costly conformational sampling steps by which designed sequences are validated [23]. This translates to a major speedup for drug design pipelines.

To address versatility, the third criterion, we ensured that CyclicCAE could be used to sample scaffolds compatible with binding to a particular target. To do this, we first ensured that CyclicCAE supports constrained conformation generation, to produce scaffolds hosting desired motifs of varying sizes. This permits functional regions of loops, proteins, or linear peptides to be presented by a designed, rigidly-folded peptide macrocycle, with immediate value in peptide therapeutic development, where stable presentation of functional motif geometry is essential. We demonstrated this functionality by using CyclicCAE to present real motifs taken from a protein loop (the N-formyl peptide receptor 2), from a linear peptide (glucagon-like peptide 1), and from an existing computationally designed peptide macrocycle (NDM1 inhibitor 1G). For motifs of 3 residues or less, CyclicCAE excels in accuracy, producing 6 to 9 residue scaffolds in which motif geometry is preserved to within 0.5 Å mainchain heavy-atom RMSD on average, while also executing very quickly, producing 1,000 candidate scaffolds in minutes on a single CPU core. Even 4-residue motifs could be generated with reasonable accuracy and speed (**Fig. 4**). In comparison, even for unconstrained generation without motifs, producing 1,000 outputs in minutes on a single core is well beyond `GeneralizedKIC`‘s abilities. In past work sampling macrocycle conformations starting from a known motif, it was generally necessary to carry out hundreds of parallel `GeneralizedKIC` runs on hundreds of CPU cores across a large computing cluster [12, 13]. In addition, by including a Lennard-Jones style repulsive term in the loss function used during inference, we ensured that CyclicCAE is able to sample and relax scaffolds in the context of a target protein binding pocket, minimizing clashes with the target while preserving favourable internal interactions like backbone hydrogen bonds that promote rigid structure (**Fig. 5**). The combination of motif presentation and pocket context-awareness ensures that CyclicCAE is as versatile and applicable to macrocycle drug design as was `GeneralizedKIC`. Combined with its proven accuracy and greater speed, this makes CyclicCAE a powerful new tool for drug discovery.

## Conclusions

This work introduces CyclicCAE, a general deep learning model for generating heterochiral macrocycle scaffolds *de novo*. Using training data taken from high-efficiency peptide macrocycle conformational sampling simulations, we demonstrate the CyclicCAE model architecture’s functionality by training it to encode and to efficiently sample low-energy conformations of 6- to 9-residue poly-glycine macrocycles. Our work demonstrates that, having exhaustively sampled the conformation space of a class of macrocycle once using a more expensive simulation-based method, our trained CyclicCAE model is able to accurately capture the energetically favourable conformations accessible to the class, while enabling much more rapid subsequent sampling of these conformations. We have also shown that CyclicCAE offers the versatility to generate scaffolds presenting a desired binding motif, and to produce scaffolds shaped to fit within binding pockets on target proteins. Significantly, nothing about the model is specific for peptides of these sizes, for poly-glycine chains, or even for *α*-amino acids. Given suitable training data (from `GeneralizedKIC`-based sampling, MD simulations, experimental databases, or other sources), this general architecture can be used for any size of macrocycle built from any combination of amino acids, peptoid monomers, or other polymeric building-blocks. We anticipate that CyclicCAE will be a powerful tool for reducing the computational cost of designing new macrocyclic therapeutics, and for carrying out rapid conformational sampling simulations to validate designs, thus accelerating macrocycle drug discovery pipelines.

## Abbreviations Used

C5AR1: C5a anaphylatoxin chemotactic receptor 1
DL: deep learning
DoF: degree of freedom
FPR2: N-formyl peptide receptor 2
GeneralizedKIC: Generalized Kine-matic Closure
GLP1: glucagon-like peptide 1
GPCR: G-protein coupled receptor
MD: molecular dynamics
ML: machine learning
MPI: Message Passing Interface
NCAA: non-canonical amino acid
NDM1: New Delhi metallo-*β*-lactamase 1
PDB: Protein Data Bank
RMSD: root mean squared deviation.

## Supplementary Information

The Supplementary Information provides additional details on CyclicCAE architecture, training, and use, as well as on Rosetta tools implemented to enable generation of training datasets.

## Acknowledgements

PH and ACP were funded through the NIH grant DP2GM146249f. PDR and VKM were funded by the Simons Foundation. The computations reported in this paper were in part performed using resources made available by the Flatiron Institute. The Flatiron Institute is a division of the Simons Foundation. ACP thanks the Center for Computational Biology at the Flatiron Institute for hospitality while a portion of this research was carried out. He also thanks S.M. Bargeen A. Turzo and Qiyao Zhu at the Flatiron Institute, and Kevin Harnden at the University of Oregon, for their helpful conversations and for providing feedback. Additionally, he is grateful to Erik Thiede in the Department of Chemistry and Chemical Biology at Cornell for the helpful talks and ideas while working on this project. Finally, the authors are grateful to the Flatiron Institute’s Scientific Computing Core, whose staff provided invaluable assistance throughout this project.

## Code and model availability

CyclicCAE model architecture, training scripts, and weights can be found on Github (https://github.com/flatironinstitute/CyclicCAEpublic); these are released under a BSD 3-clause free and open-source licence. Current maintainers and questions should be directed through issues on Github and/or through email to Andrew Powers, Parisa Hosseinzadeh, and Vikram K. Mulligan.

## Statement on generative AI

Aside from CyclicCAE itself, generative AI was not used in the preparation of this manuscript’s text or figures.

## Extended Data Figures

**Ext. Data Fig. 1.**
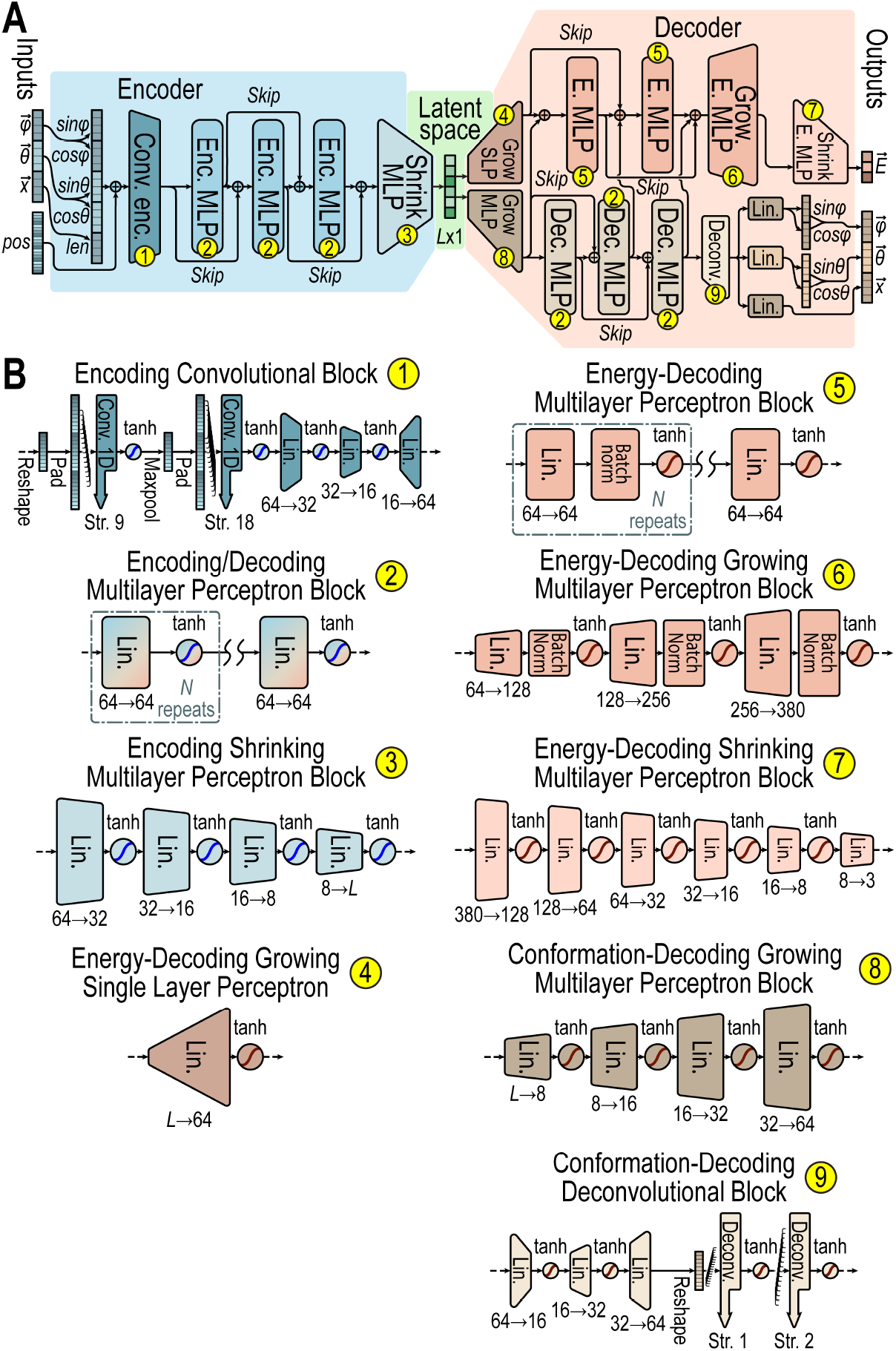
Details of CyclicCAE neural network architecture. (**A**) Overall architecture, reproduced from **Fig. 1**. Blue: encoding layers. Green: the *L*-by-1 latent space. *L* was 6 in all CyclicCAE instances trained for this work. Pink: energy-decoding layers. Beige: conformation-decoding layers. Yellow circles label blocks. The model maps an input vector of mainchain dihedrals 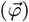, mainchain bond angles 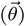, and mainchain bond lengths 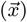, plus positional encodings (*pos*), to an output vector of energies 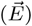, mainchain dihedrals 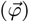, mainchain bond angles 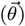, and mainchain bond lengths 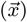. (**B**) Architectural details of blocks marked in panel **A**. Lin: linear transformation layers (with input and output dimensions indicated). Conv. 1D: one-dimensional convolutional layers, with stride indicated. Deconv.: one-dimensional deconvolutional layers, with stride indicated.

**Ext. Data Fig. 2.**
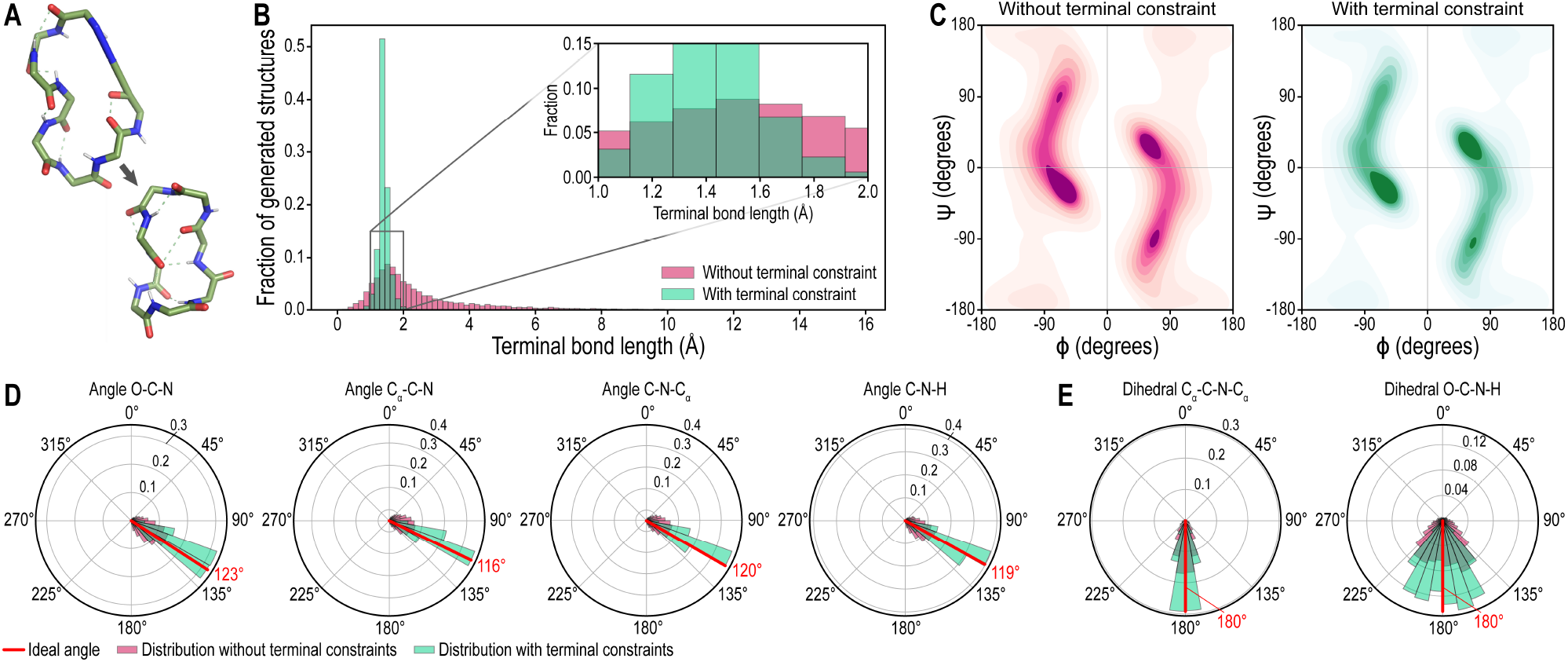
Effect of including terminal bond constraint terms (Supplementary Information section S3.2.2) during latent-space gradient descent refinement. (**A**) Representative example of impact on peptide geometry: a distorted terminal amide bond is converted into an amide bond with appropriate bond length, bond angles, and planarity (dihedral angle), while preserving other favourable structural features. (**B**) Histogram of terminal bond lengths produced from 10,000 outputs with (green) and without (magenta) the terminal bond constraint. (**C-D**) Density plot of sampled *ϕ* and *ψ* angles in 10,000 outputs, demonstrating that the distribution of torsion angles sampled from CyclicCAE is similar without (**C**, magenta) and with (**D**, green) the terminal constraint, meaning there are no unnatural effects placed on the mainchain torsions. (**D**) Polar histograms showing the distribution of terminal bond bond angles O-C-N, C_*α,i*_-C-N, C-N-C_*α,i*+1_, and C-N-H without (magenta) and with (green) the terminal constraint. Ideal bond angles are shown in red. (**D**) Polar histograms showing the distribution of terminal bond dihedral angles C_*α,i*_-C-N-C_*α,i*+1_ and O-C-N-H without (magenta) and with (green) the terminal constraint.

**Ext. Data Fig. 3.**
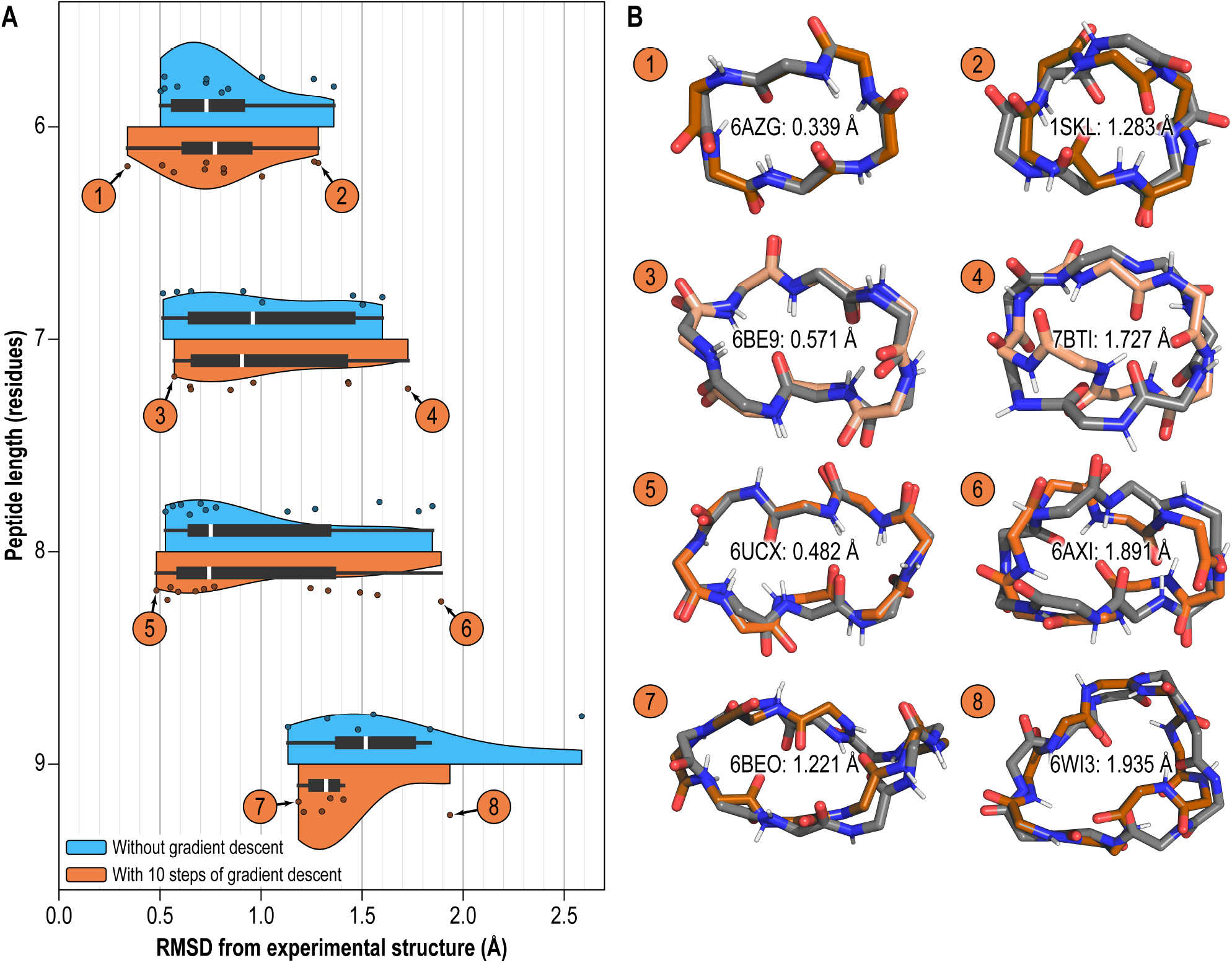
Effect of gradient descent minimization of estimated energy on reproduction of experimentally-determined peptide macrocycle conformations. (**A**) Distribution of mainchain heavy-atom RMSD values between input macrocycle conformations from PDB structures and reconstructed conformations produced by CyclicCAE, without (cyan) and with (orange) gradient descent refinement. In the latter case, 10 steps of gradient descent refinement were performed in the latent space to minimize the decoder limb’s estimate of the conformational energy. (**B**) Experimentally-determined structures (grey) and conformations produced after CyclicCAE encoding, decoding, and energymiminization steps (orage). Numbers correspond to points in panel **A**. PDB IDs and RMSD values are indicated.

**Ext. Data Fig. 4.**
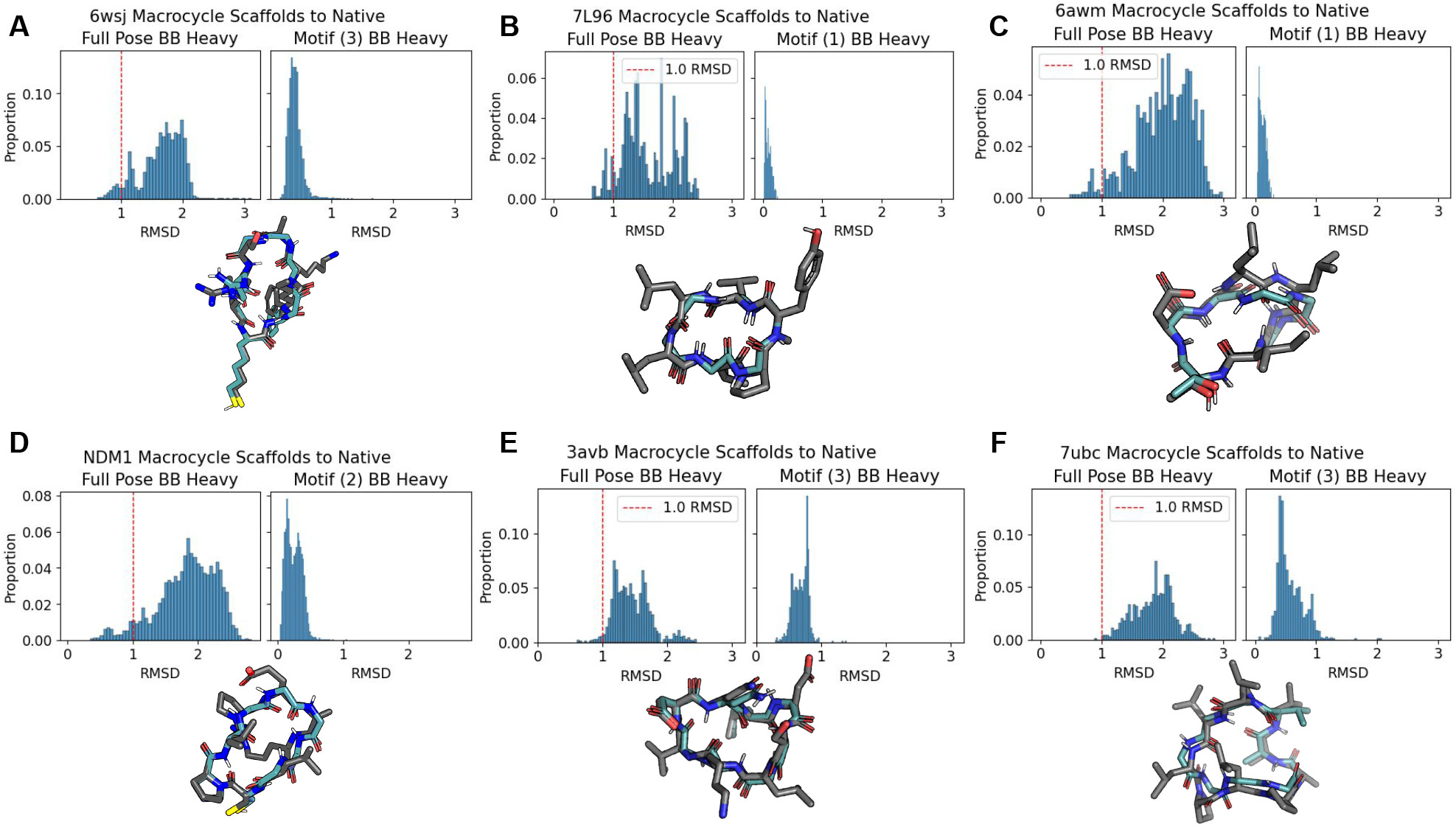
Reproduction of single-residue motifs from known cyclic peptides. 1,000 macrocycle conformations were sampled that could present a specific motif taken from six peptide macrocycles found in the PDB. (**A-F**) In each panel, the left graph shows the mainchain heavy-atom RMSD of the scaffolds generated by CyclicCAE to the native PDB structure. The right graph is the RMSD of the motif mainchain heavy-atoms in the CyclicCAE-generated scaffolds to the motif mainchain heavyatoms in the native PDB structure. The index of the single-residue motif is shown in round brackets in the graph title. Sticks representations show the native PDB structure in grey and the lowest-RMSD CyclicCAE-generated scaffold overlain. CyclicCAE is consistently able to reproduce the original peptide conformations, as well as to propose diverse new scaffolds that can also present the motif.

## Supplementary Information

### S1 Existing Rosetta methods used in this work

#### S1.1 Obtaining and compiling Rosetta

The work described here relied on the Rosetta software suite [1, 2]. Although not open-source, Rosetta is made available for free for non-commercial use from https://github.com/RosettaCommons/rosetta, and is licenced for commercial use through UW Comotion (see https://els2.comotion.uw.edu/product/rosetta).

Instructions for compiling Rosetta are available at https://docs.rosettacommons.org/demos/latest/tutorials/install_build/install_build. Note that if you are processing a large number of tasks, we recommend building Rosetta with support for parallelism through the Message Passing Interface (MPI) and through POSIX threads. This may be achieved by adding `extras=cxx11thread,mpi,serialization` to the `scons` compilation command. Alternatively, if you are using the Masala-patched version of Rosetta (see [3]), you may use `extras=masala,mpi,serialization`, which will also add multi-threading support. MPI support will allow automatic distribution of design jobs over available CPU cores, on a single node or across nodes on a network, and multi-threading support adds additional shared-memory parallelism within jobs at key steps such as interaction graph precomputation. This is particularly strongly recommended for mainchain conformational sampling for validation of designs with Rosetta’s `simple_cycpep_predict` application (see section S1.3, since the MPI-parallelized version carries out additional parallelized analysis, such as automatic computation of fold propensity (*P*_near_) and estimation of Δ*G*_*folding*_.

Some of the protocols, such as the `GeneralizedKIC` sampling script described in section S1.2.1, require the `PyRosetta` scripting language. For instructions on obtaining and building `PyRosetta`, see https://docs.rosettacommons.org/docs/latest/scripting_documentation/PyRosetta/PyRosetta.

#### S1.2 Generalized Kinematic Closure (GeneralizedKIC) for macrocycle conformational sampling

Generalized Kinematic Closure, or `GeneralizedKIC`, was the prior state of the art method for generating heterochiral macrocycle poly-glycine scaffolds *in silico*, described in references [4, 5, 6, 7] and reviewed in detail in [8]. It uses a robotics-inspired algorithm to very rapidly solve kinematic equations to sample conformations of a chain of atoms with a fixed rigid-body transform between the chain’s start and end [9, 10]. This is coupled with filtering to discard closure solutions with undesired properties. Since many solutions may be discarded that have, for instance, too few mainchain hydrogen bonds or too many residues in unfavourable regions of Ramachandran space, `GeneralizedKIC` can require many closure attempts to find energetically favourable solutions.

##### S1.2.1 PyRosetta script for GeneralizedKIC-based mainchain sampling

To compare CyclicCAE to the previous state of the art, we used `GeneralizedKIC` to sample an ensemble of conformations of peptide macrocycles ranging from 6 to 9 amino acid residues. We wrote a custom `PyRosetta` script to build our conformational ensembles, which is available as *inputs/insolution genkic*.*py* in the CyclicCAE git repository (see section S4.1).

The script starts by building the peptide macrocycle geometry. When generating conformational ensembles using `GeneralizedKIC`, a poly-glycine linear pose, if heterochiral structures are desired, is built using Rosetta. A Rosetta `FoldTree` is set to designate the central residue of the `Pose` (*i*.*e*. Rosetta structure) as the root atom to minimize lever arm effects. The script next removes terminal types, adds cutpoint variants (`CUTPOINT_LOWER` and `CUTPOINT_UPPER`), and declares a chemical bond connecting the N_1_ and C_*N*_ atoms.

The script then randomizes the *ϕ* and *ψ* dihedral angles of an central residue (which will serve as the “anchor” residue for kinematic closure), using the Rosetta `RandomizeByRamaPreProMover` to ensure that these angles are drawn from the probability distribution of accessible angles given the Ramachandran map of glycine. It then configures a Rosetta `GeneralizedKIC Mover` object, setting 500 closure attempts, a minimum of 1 final solution, selection of the top solution using the `lowest_energy_selector`, and `ref2015` as the energy function. Three “pivot” residues are chosen, with two adjacent to the “anchor” residue and one randomly selected from the remaining residues. These residues’ *ϕ* and *ψ* values will be calculated analytically to ensure loop closure. All non-pivot residues are set to be perturbed by the `GeneralizedKIC randomize_backbone_by _rama _prepro` perturber, which randomizes mainchain *ϕ* and *ψ* dihedral angles biased by the Ramachandran distribution for glycine (or the amino acid type in question).

Prior to selecting a top solution for each repeat, the script discards solutions with oversaturated hydrogen bond acceptors (using the Rosetta `OversaturatedHbondAcceptorFilter` with an `hbond_energy_cutoff` value of -0.25). This process is repeated until the desired number of structures (specified with the `--nstruct` option) has been obtained.

Please refer to the `README.md` file for full instructions on running this script. A sample commandline is shown in Supplementary Listing S1.

**Supplementary Listing S1.**
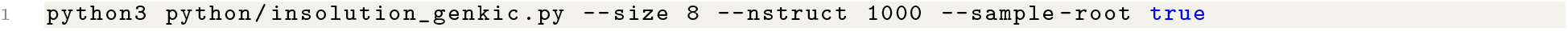
Example commandline command for running the `GeneralizedKIC` Python script to produce 1,000 8-mer backbones.

#### S1.3 The simple cycpep predict application for parallelized macrocycle conformational sampling

`GeneralizedKIC` is written to be a module usable in larger Rosetta protocols, such as those scripted with `RosettaScripts` [11] or `PyRosetta` [12] (see section S1.2.1). Additionally, Rosetta’s `simple cycpep predict` application [5] wraps `GeneralizedKIC`, and provides a convenient way of sampling large numbers of macrocycle conformations on a large computing cluster, with parallelization provided by the Message Passing Interface (MPI). We used `simple cycpep predict` to sample large numbers of conformations of poly-glycine macrocycles, ranging from 6 to 9 residues, in order to generate training data for CyclicCAE.

##### S1.3.1 Using simple cycpep predict to sample poly-glycine macrocycle conformations to produce training data

To sample large numbers of conformations of poly-glycine macrocycles using `simple_cycpep_predict` in order to assemble training datasets for CyclicCAE, it is first necessary to obtain and compile Rosetta as described in section S1.1, using the MPI and multi-threaded, or MPI and Masala, compilation options. (Single-process, singlethreaded execution of `simple_cycpep_predict` would be unable to sample sufficient conformations in a reasonable amount of time.)

Next, it is necessary to prepare a working directory, with an `inputs/` sub-directory. To the `inputs/` directory should be added the following file, containing commandline flags for the `simple_cycpep_predict` application:

**Supplementary Listing S2.**
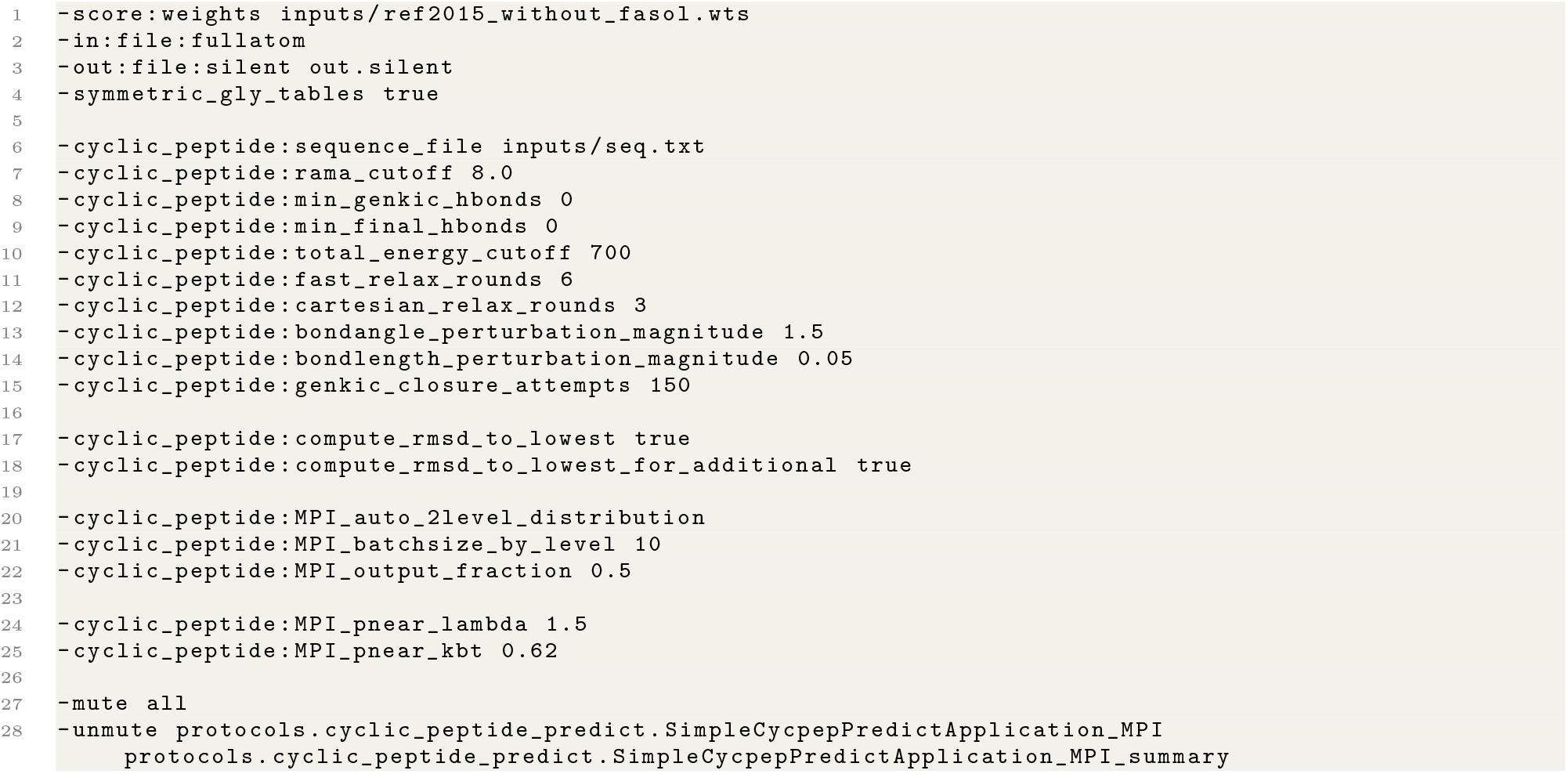
File `inputs/rosetta.flags` containing commandline flags for the `simple cycpep predict` application.

Next, it is necessary to define the scoring function that will be used during relaxation and energy calculation steps of the sampling protocol. We use a modified version of Rosetta’s default `ref2015` energy function, with the implicit solvation terms deactivated. This should be specified in a `inputs/ref2015 without fasol.wts` file, containing the lines shown in Supplementary Listing S3:

**Supplementary Listing S3.**
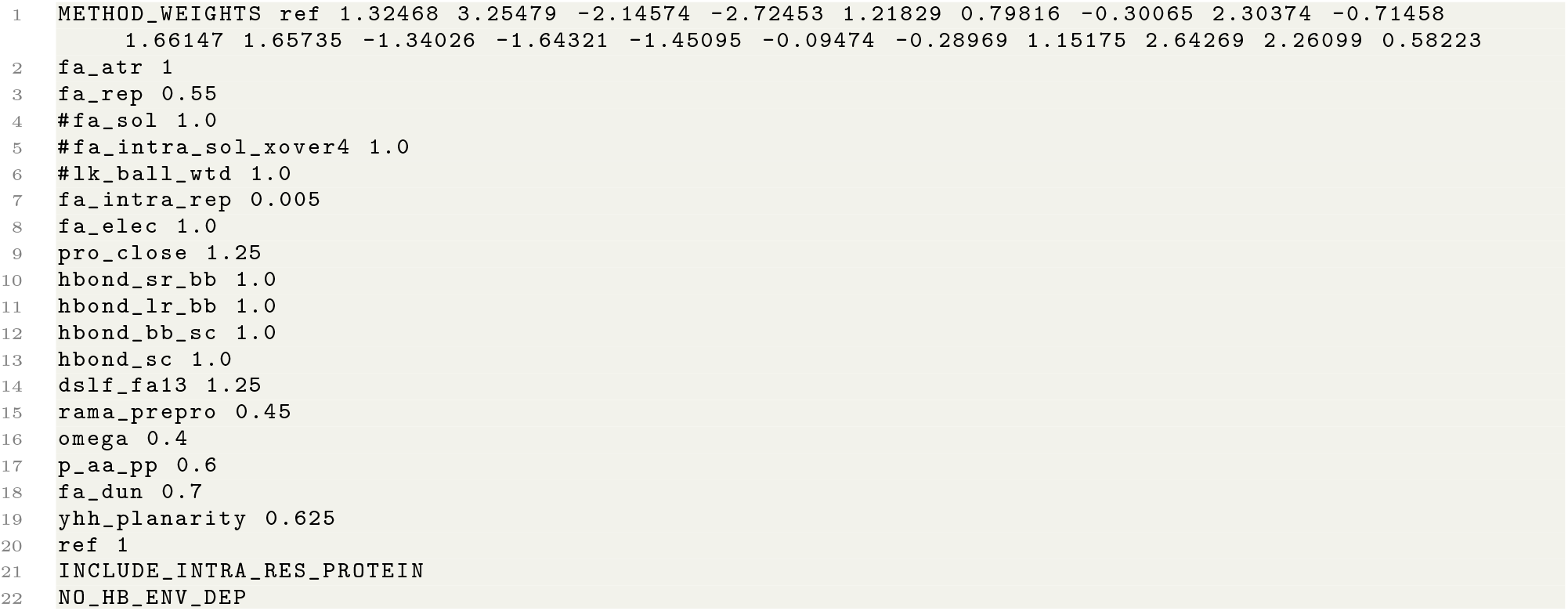
File `inputs/ref2015 without fasol.wts`, defining the scoring function used for relaxation and scoring during poly-glycine conformational sampling.

We also need to specify the sequence of the macrocycle that we will be sampling, in a `inputs/seq.txt` file:

**Supplementary Listing S4.**
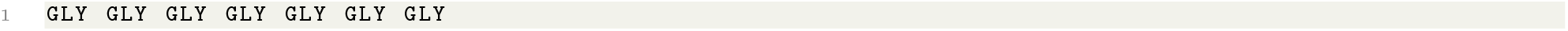
File `inputs/seq.txt`, specifying the sequence to sample.

In Supplementary Listing S4, a 7-residue poly-glycine sequence has been specified. Longer sequences may be specified by adding additional instances of “GLY”.

Finally, we need the command that runs `simple cycpep predict`:

**Supplementary Listing S5.**
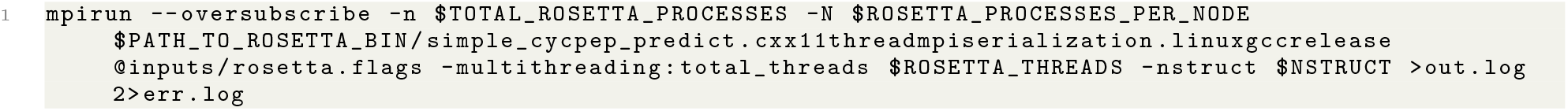
The `simple cycpep predict` execution command.

In Supplementary Listing S5, above, it is assumed that certain environment variables have been set. These are described in Supplementary Table S1.

##### S1.3.2 Modifications to simple_cycpep_predict inputs when used with RosettaQM

`RosettaQM` is a recently-developed bridge between the Rosetta software suite and quantum chemistry packages such as `GAMESS` [13] or `ORCA` [14]. It permits quantum chemistry calculations in the context of any Rosetta protocol. In `simple_cycpep_predict`, RosettaQM may be used for quantum chemistry-based geometry optimization (relaxation), and for quantum chemistry-based calculation of conformational energies. We used this to add additional energy terms to our training data, which CyclicCAE was trained to predict alongside the Rosetta `ref2015` energy. Note that RosettaQM will be released shortly, and described in a separate publication (manuscript in preparation). Until its release, please contact the authors for access to the software if you are interested in reproducing this part of our protocol. CyclicCAE may also be trained or used without RosettaQM (*i*.*e*. using training data containing only energies computed with Rosetta’s energy functions).

To use `simple_cycpep_predict` with `RosettaQM`, it is first necessary to apply the `RosettaQM` patch to Rosetta and to recompile. Once this is done, the next step is to add lines to the flags file specifying configuration files for quantum chemistry-based geometry optimization steps and energy calculation steps:

**Supplementary Listing S6.**
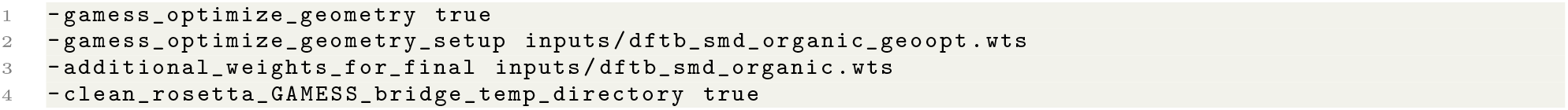
Additional commandline flags added to `inputs/rosetta.flags` to enable quantum chemistry-based geometry relaxation and energy calculation for each sample, for use with the `RosettaQM` extension.

In Supplementary Listing S6, these four flags specify that `simple_cycpep_predict` should perform quantum chemistry-based geometry optimization following force field-based relaxation of each sampled macrocycle conformation. They specify a scoring function configuration file for a quantum chemistry-based geometry optimization step and for an additional final quantum chemistry-based energy calculation step. The final line specifies that temporary files written during these quantum chemistry steps should be deleted rather than retained.

**Supplementary Table S1.**
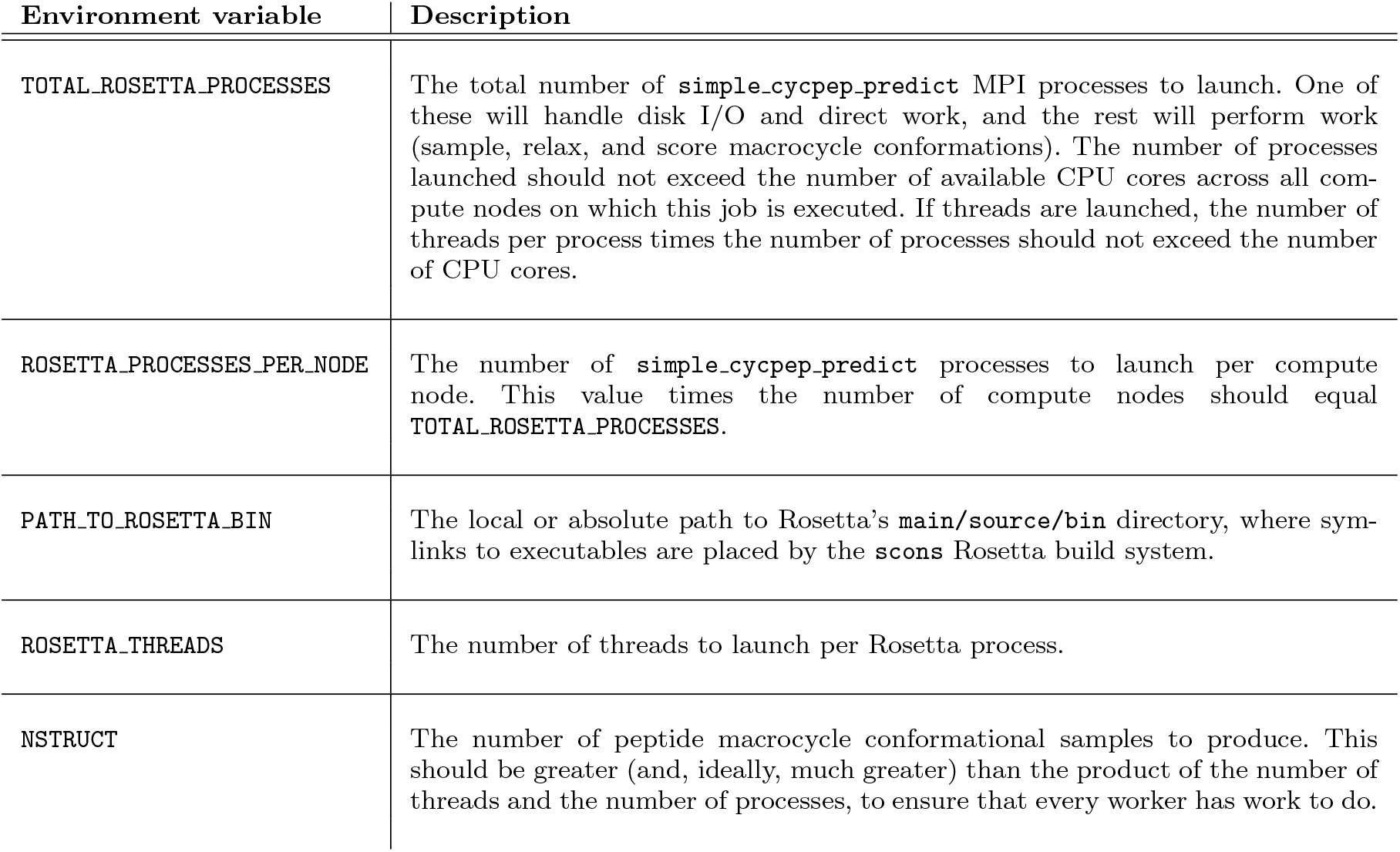
Environment variables that must be set to run simple cycpep predict using the command shown in Supplementary Listing S5.

The contents of the `inputs/dftb_smd_organic_geoopt.wts` and `inputs/dftb _smd _organic.wts` files are shown in Supplementary Listings S7 and S8, respectively. In this case, these specify that the Density Functional Tight Binding (DFTB) semi-empirical quantum chemistry method should be used for both geometry optimization and energy calculation, though any semi-empirical, DFT, or Møller–Plesset method supported by `GAMESS` or `ORCA` could be used (including Fragment Molecular Orbital methods supported by `GAMESS` [15]). The user controls the tradeoff between speed and accuracy.

**Supplementary Listing S7.**
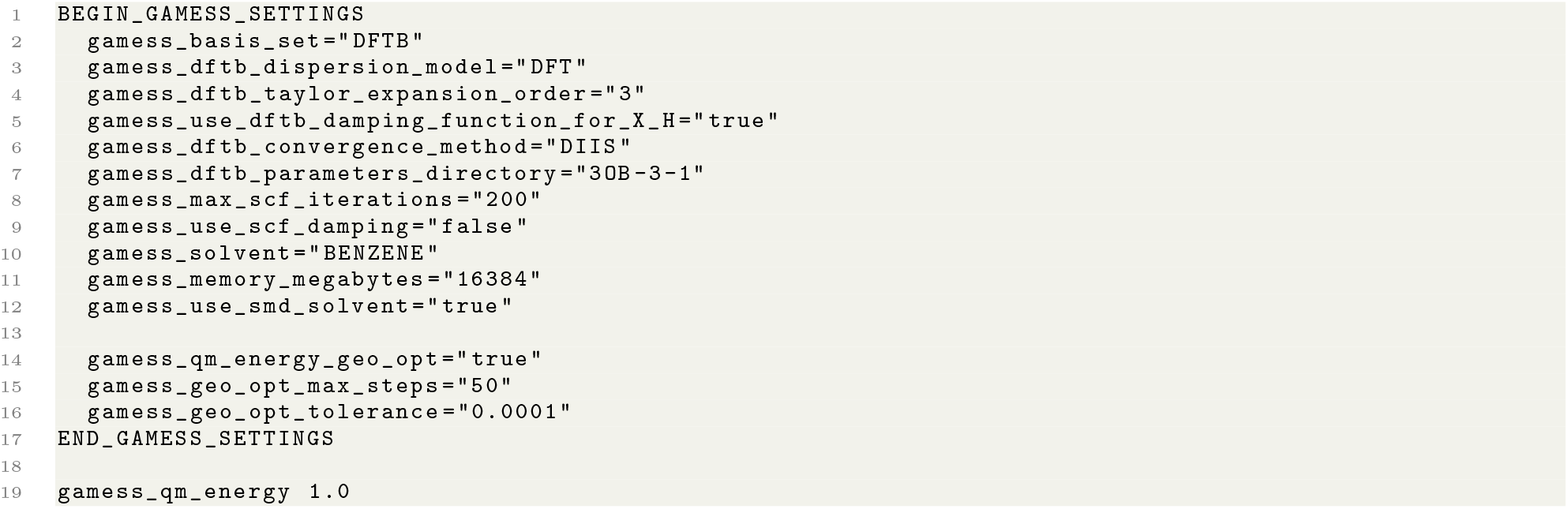
File `inputs/dftb_smd_organic_geoopt.wts`, used to configure quantum chemistry-based relaxation of macrocycle geometries using the `RosettaQM` extension and the `GAMESS`quantum chemistry software.

**Supplementary Listing S8.**
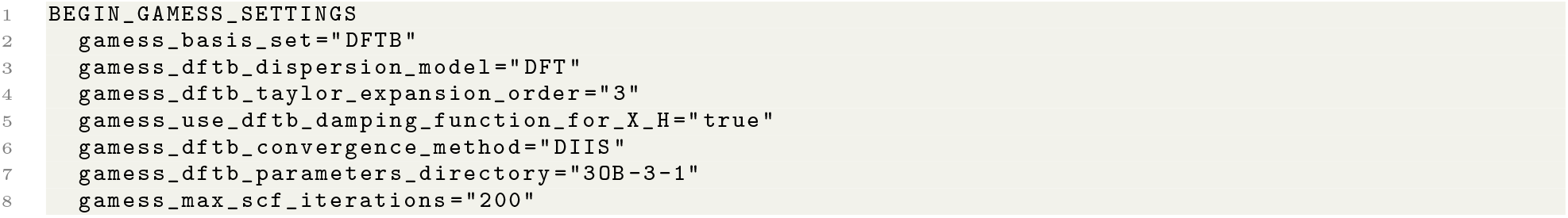
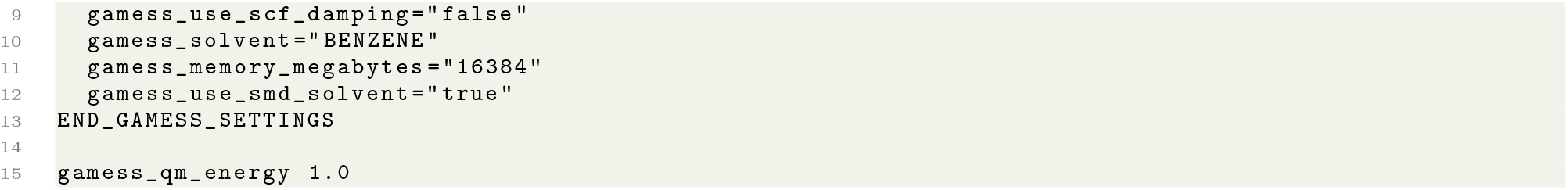
File `inputs/dftb_ smd_ organic.wts`, used to configure quantum chemistry-based calculation of macrocycle conformational energies using the `RosettaQM` extension and the `GAMESS` quantum chemistry software.

Finally, the command to run `simple_ cycpep_ predict` must be modified with additional inputs specifying where the `GAMESS` (or `ORCA`) executable is, and where temporary output should be written. The updated command is shown in Supplementary Listing S9 The additional environment variables that should be set are listed in Supplementary Table S2.

**Supplementary Listing S9.**
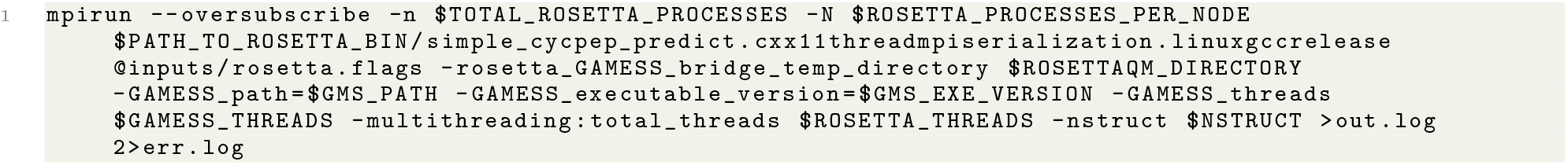
Modifications to the `simple_ cycpep_ predict` execution command when running with the `RosettaQM` extension and `GAMESS` quantum chemistry calculations.

##### S1.3.3 Using simple cycpep predict to validate designed peptide macrocycles

Peptide designs produced using CyclicCAE- or `GeneralizedKIC`-generated scaffolds and the `FastDesign`-based design script described in section S1.5.1 were computationally validated using the Rosetta `simple _cycpep_ predict` application to sample conformations of the designed amino acid sequence and to determine the extent to which the sequence favoured folding into the desired conformation. The application also calculated the *P*_near_ metric [16], defined in Eq. (S1).

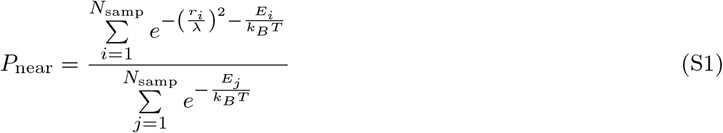

**Supplementary Table S2.**
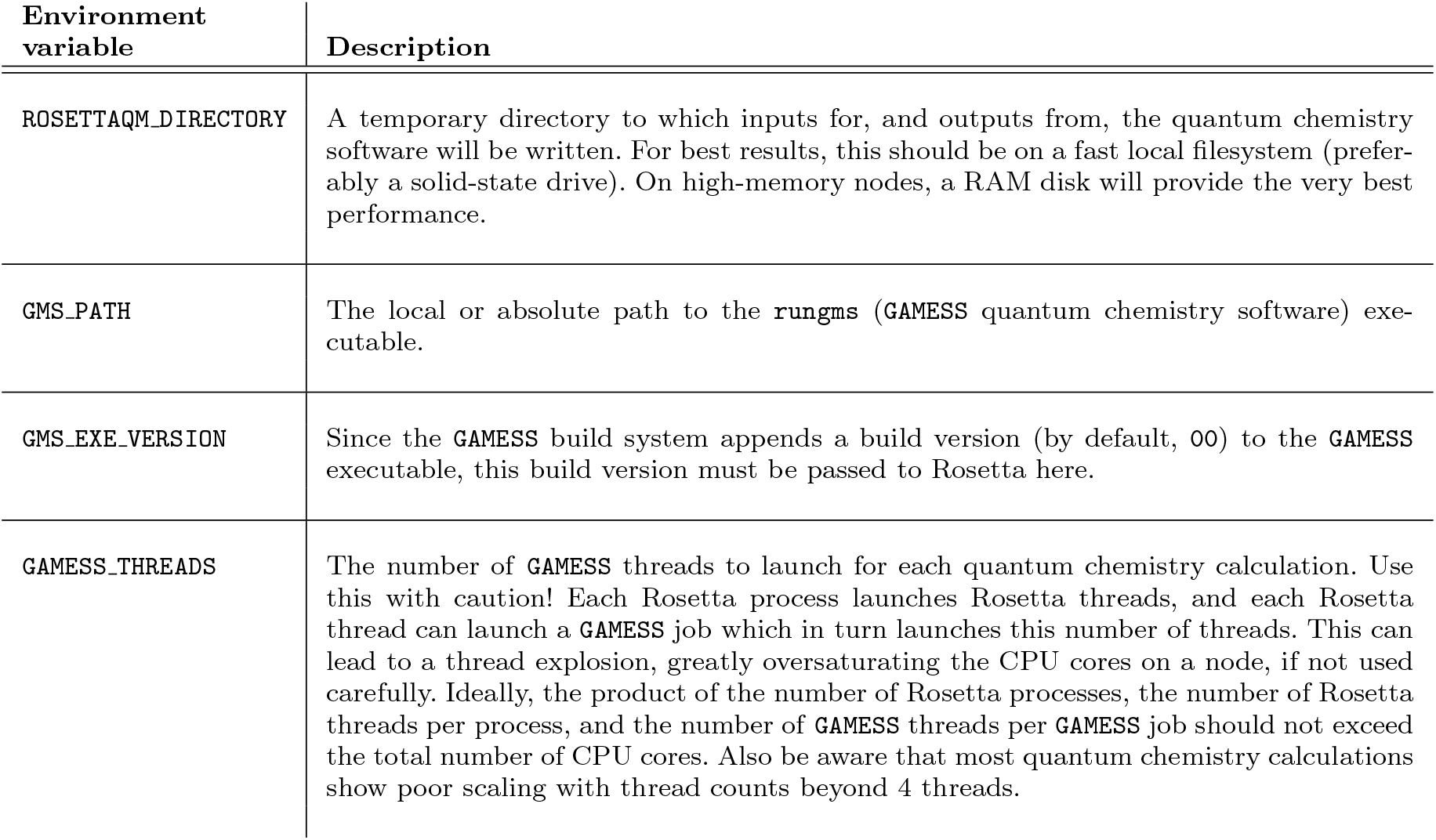
Additional environment variables that must be set to run simple_ cycpep _predict with RosettaQM, using the command shown in Supplementary Listing S9.

In Eq. (S1), *k*_*B*_ is the Boltzmann constant, *N*_samp_ is the number of sampled conformations, *r*_*i*_ is the root mean squared deviation (RMSD) of the *i*_th_ sample from a desired conformation, and *E*_*i*_ is the energy of the *i*_th_ sample. The user-set parameters *λ* and *T*, measured in Ångstroms and Kelvin, respectively, govern the RMSD proximity that a sample must have to be considered “native-like”, and the extent to which increases in energy decrease the probability of a sample, respectively. For practical reasons, the `simple_cycpep_predict` application accepts a value of *k*_*B*_*T* in 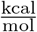 instead of a value of *T* in Kelvin. For our purposes, we set *λ* = 1.5 Å (suitable for small macrocycles) and 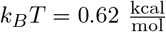 (corresponding to physiological temperature).

To validate designed cyclic peptides, the protocol described in section S1.3.1 was modified as follows. First, the sequence file shown in Supplementary Listing S4 was replaced with one listing the designed amino acid sequence. Rosetta “basenames” (*e*.*g*. “ALA” for l-alanine, “DALA” for d-alanine) must be included in this file. Second the designed structure was provided by adding `-in:file:native design.pdb` to the `rosetta.flags` file shown in Supplementary Listing S2.

#### S1.4 Using the energy based clustering application to cluster sampled macrocycle mainchain conformations

We used Rosetta’s `energy_ based _clustering` application to cluster the sampled macrocycle mainchain conformations in order to produce smaller datasets for use during the early stages of training, to speed up these early training steps. The `energy based clustering` application uses a simple “cookie cutter” algorithm for clustering: it identifies the lowest-energy sample, marks that as the centre of the first cluster, computes a vector of RMSD values between this sample and all other samples, and assigns all samples within a threshold distance to the first cluster. It then removes these samples from the pool of unassigned samples and repeats the steps above to assign the second, third, fourth, *etc*. clusters, until there are no more samples left to assign to clusters. Documentation for the `energy_ based _clustering` application is available from https://docs.rosettacommons.org/docs/latest/applicationdocumentation/analysis/energy-based_clustering_application.

We ran the `energy_ based _clustering` application with the options shown in Supplementary Listing S10:

**Supplementary Listing S10.**
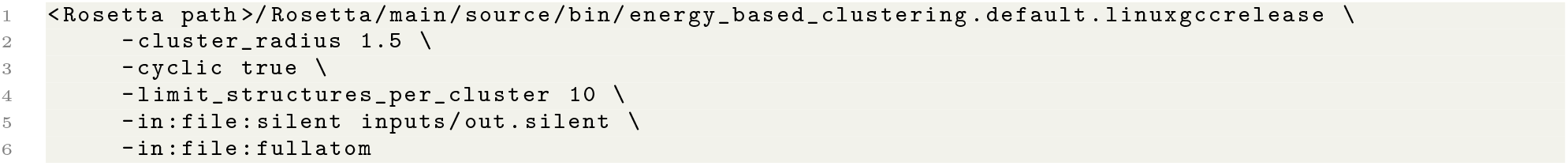
Commandline command for running `energy_ based _clustering`.

In Supplementary Listing S10, we assume that the output Rosetta silent file (containing sampled conformations) from `simple cycpep predict` sampling is placed in an `inputs/` directory. Output files are written to the working directory, with only the lowest-energy ten structures per cluster written to disk. This served as our reduced training dataset for early training steps, as described in section S4.3.

#### S1.5 Macrocycle design tools

The Rosetta software suite’s tools for designing peptide sequences have been described elsewhere [4, 5, 7, 17, 16]. Briefly, Rosetta uses a high-efficiency simulated annealing-based algorithm, called the `Packer`, to search through combinatorial possibilities of amino acid residue identities and sidechain conformation, collectively called *rotamers* (see [18] for a detailed review). Its `FastDesign` module alternates calls to the `Packer` with gradient descent relaxation steps, performed by the Rosetta `Minimizer` [4, 1]. During both sequence design and relaxation, the objective function minimized is typically an energy function such as the `ref2015` energy function [19]. This may be augmented with design-centric guidance terms such as `aa _composition` to control overall amino acid composition, `voids penalty` to promote designs with well-packed cores, or other terms to penalize undesired features or encourage desired features in designs [8].

##### S1.5.1 RosettaScripts design script

With both the set of peptide macrocycles scaffolds produced by CyclicCAE and that produced with `GeneralizedKIC`, we used a `FastDesign`-based protocol, controlled with the `aa composition` design-centric guidance term appended to the `ref2015` energy function [5], to design peptide sequences to try to maximize fold propensity. `RosettaScripts` scripts for designing a peptide sequence given a fixed mainchain sampled with either `GeneralizedKIC` or CyclicCAE are provided in the CyclicCAE Github repository (https://github.com/flatironinstitute/CyclicCAE_public) in the `xml/` sub-directory. These are summarized in Supplementary Table S3.

Users may run this script as follows:

1. Obtain and compile Rosetta as described in section S1.1.
2. Place input mainchain PDB files in an appropriate directory. For our purposes, we will assume that they are in a `pdbs/` sub-directory of a working directory into which outputs will be written.
3. Symlink the CyclicCAE `xml/` sub-directory. This is necessary to allow the script to find the other files that it needs, since they are specified relative to the working directory. On Linux, this can be done as follows:

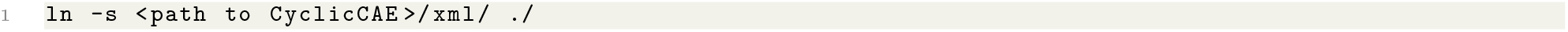

3 Run Rosetta. If one is using the default build, the command will be:

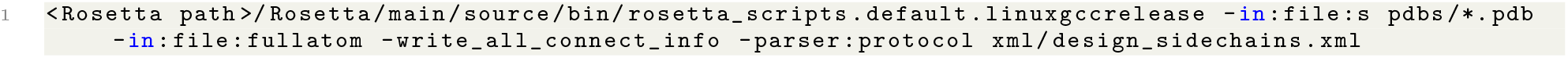

In the above, `linuxgccrelease` should be replaced with `macosclangrelease` on Macintosh systems.

If one is using the MPI build, the command will vary, depending on one’s MPI flavour and cluster configuration. A possible execution command would be:

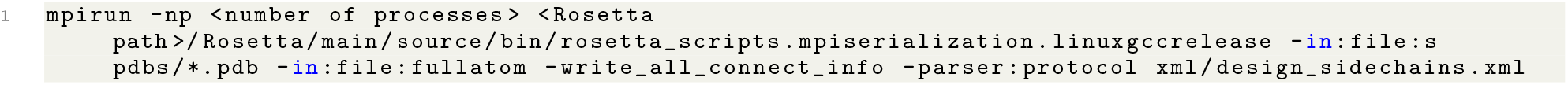

Additionally, one may add the commandline option `-mute all` to reduce the verbosity of tracer output, and may pipe tracer output to output and error logfiles by appending `>out.log 2>err.log`.

**Supplementary Table S3.** Files located in the `xml/` sub-directory of the CyclicCAE Github repository (https://github.com/flatironinstitute/CyclicCAE_public), which permit sequence design using Rosetta given an input set of peptide macrocycle mainchains sampled with CyclicCAE, `GeneralizedKIC`, or another sampling method.

| Filename | Description |
| --- | --- |
| `design_sidechains.xml` | A RosettaScripts XML file which takes as input one or more PDB files representing cyclic mainchains sampled with CyclicCAE or another sampling method, and produces as output sequences and structures of the relaxed designs. The sequence of steps is: (a) declaration of the N-to-C peptide bond, and addition of appropriate Rosetta constraints to ensure that cyclic geometry is preserved during relaxation, (b) addition of composition constraints from file `desired_makeup.comp`, (c) mutation of all positions compatible with L-proline to L-proline, and all positions compatible with D-proline to D-proline, (d) addition of biases for design to favour the D- or L-proline residue identities that have been added, but to permit mutation to other residue types if they are strongly favoured, (e) execution of Rosetta’s **FastDesign** protocol to perform alternating rounds of sidechain optimization and all-dihedral relaxation, and (f) execution of Rosetta’s **FastRelax** protocol to carry out additional relaxation. |
| `d_res.resfile` | A Rosetta resfile indicating that design should be limited to the D-amino acid mirror-images of the 19 canonical L-amino acids, with the exception of D-cysteine. This is used at positions in the peptide where the $\phi$ mainchain torsion angle is positive. Invoked by `design_sidechains.xml`. |
| `l_res.resfile` | A Rosetta resfile indicating that design should be limited to the 19 canonical L-amino acids, with the exception of L-cysteine. This is used at positions in the peptide where the $\phi$ mainchain torsion angle is negative. Invoked by `design_sidechains.xml`. |
| `desired_makeup.comp` | A Rosetta amino acid composition file used to control the amino acid composition during design, to promote sequences that stabilize rigid macrocycle folds. This file specifies that (a) sequences with fewer than 2 D- or L-proline residues should be heavily penalized, (b) sequences should be penalized if they do not contain at least one L-aspartate or L-glutamate residue, to permit sidechain tethering to the resin during synthesis (a common design constraint to permit on-resin cyclization), (c) sequences should be penalized for containing fewer than 1 D- or L- arginine or lysine residue (to promote solubility), (d) sequences should be penalized if they contain more than 25% D- or L- arginine or glutamate (to avoid difficult-to-synthesize sequences), and (e) sequences should be penalized if they contain no hydrophobic residues. This serves to guide the Rosetta packer towards sequences that have desired properties. Invoked by `design_sidechains.xml`. |

**Supplementary Table S4.** Number of peptide macrocycle designs (from a pool of 5,000 produced for benchmarking) that passed design filters, by macrocycle size and scaffold generation method.

| Macrocycle size<br>(residues) | GeneralizedKIC based<br>designs passing<br>filters | CyclicCAE based<br>designs passing<br>filters |
| --- | --- | --- |
| 6 | 4,111 | 4,376 |
| 7 | 4,064 | 4,627 |
| 8 | 4,272 | 4,621 |
| 9 | 4,531 | 4,624 |

### S2 New Rosetta methods developed for this work

#### S2.1 Rosetta patch availability and installation

Although CyclicCAE is a standalone neural network, with trained weights for 7 to 9 residue peptides provided in the CyclicCAE Git repository, various Rosetta modifications were needed to facilitate the generation of training data, or the visualization of CyclicCAE-generated structures. These are made available as a patch for Rosetta 3.15. To apply the patch, copy it to the `Rosetta/main` directory. Then navigate to this directory and run the command shown in Supplementary Listing S11. This has been tested with Rosetta Git SHA1 `bbb6a2d27c70be05b2fb2409f3920b5894967d60`.

**Supplementary Listing S11.**
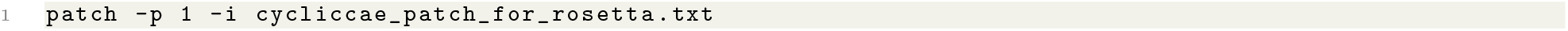
Commandline inputs to patch Rosetta.

The effect of the patch is to add the `extract_dof_vectors` application used for generating training data (described in section S2.2), the `dof_ vectors_ to _poses` application used to visualize CyclicCAE output (described in section S2.3), and the `align _ml _backbones_ to _original _target` application used to place CyclicCAE-generated scaffolds in a binding pocket on a target during binder design (described in section S2.4). The support class `DoFSummary` is also added by the patch, as described in section S2.2.

The patch also makes modifications to `Rosetta/main/source/src/basic/options/options rosetta.py` to add the `conformational_ autoencoder` options group, which contains commandline options for the `extract_ dof _vectors` and `dof _vectors_ to _poses` applications. It also makes small adjustments to the `PDBPoseInputStream` class, defined in `Rosetta/main/source/src/core/import pose/pose stream/PDBPoseInputStream.cc`, and to the `SilentFilePoseInputStream` class, defined in `Rosetta/main/source/src/core/import pose/posestream/SilentFilePoseInputStream.cc`, to permit more efficient processing of many structures.

The Rosetta patch is located in `rosetta _patch/cycliccae _patch _for _rosetta.txt` in the CyclicCAE Git repository (https://github.com/flatironinstitute/CyclicCAE_public). It is released under a free and open-source BSD 3-clause licence.

#### S2.2 A Rosetta application for generating training data extract dof vectors

We wrote a new Rosetta application called `extract dof vectors`, intended to take as input sets of conformational samples for a macrocycle of a given size, and to produce as output compact binary-format records of the vectors of salient degrees of freedom from the structures, which in turn could be used for training CyclicCAE. This application also generates a matrix of RMSD values (which is also needed for training) and writes this in binary format. After patching Rosetta, source code for this application may be found in `Rosetta/main/source/src/apps/public/cyclic peptide/extract dof vectors.cc`. This makes use of the `DofSummary` class added by the patch to `Rosetta/main/source/src/protocols/conformational autoencoder/DofSummary.cc`.

To use the `extract _dof _vectors` application, first, install the patch described in section S2.1. Next, compile all Rosetta applications, preferably with either the `extras=cxx11thread` or `extras=masala` option added to the Scons command to enable multi-threading – *e*.*g*. `python3 ./scons.py mode=release extras=cxx11thread bin`. Once this has been done, the application may be run as shown in Supplementary Listing S12.

**Supplementary Listing S12.**
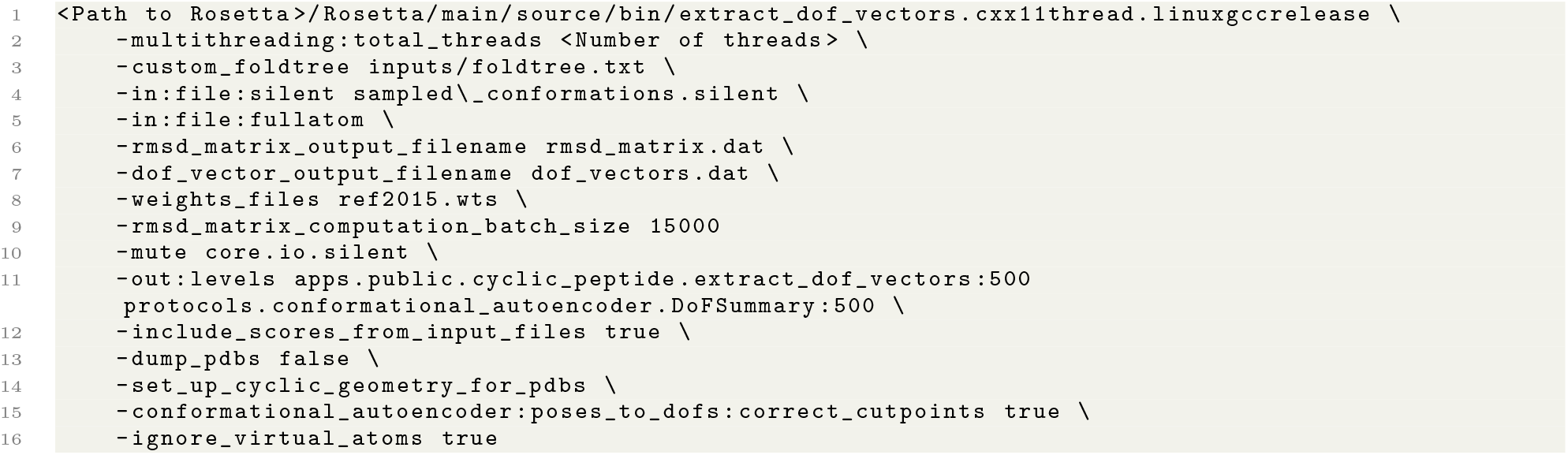
Commandline inputs to extract degree-of-freedom information and to generate an all-to-all RMSD matrix for a set of conformations sampled previously and found in `sampled conformations.silent`.

In Supplementary Listing S12, `cxx11thread` should be replaced with `masala` if the Masala build of Rosetta (see [3]) is used, and `linuxgccrelease` should be replaced with the user’s operating system and compiler as needed – for instance, `macosclangrelease`. We strongly recommend parallelizing this computation over all available CPU cores by setting `-multithreading:total_ threads` to the number of cores available on the node; this will greatly accelerate the RMSD matrix calculation. For information about all commandline options, the `extract_ dof_ vectors` application may be run with the `--help` flag. The use of `extract _dof _vectors` in processing training data for this study is further described in section S3.1.

#### S2.3 A Rosetta application for visualizing CyclicCAE conformations produced as output: dof vectors to poses

We also wrote a Rosetta application for converting the output from CyclicCAE back into Rosetta Poses, and for writing these to disk in Rosetta silent file format. This application, called `dof_vectors_ to _poses`, is found in `Rosetta/main/source/src/apps/public/cyclic _peptide/dof _vectors_ to _poses.cc` after patching Rosetta as described in section S2.1. After compiling, and given output from CyclicCAE in `cycliccae _output.dat`, the application may be executed as shown in Supplementary Listing S13 to produce `all atom structures.silent`. This may in turn be converted to a set of PDB files using Rosetta’s `extract pdbs` application (see https://docs.rosettacommons.org/docs/latest/extract-pdbs).

**Supplementary Listing S13.**
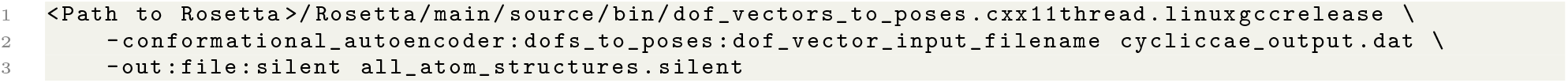
Commandline inputs to convert degree-of-freedom vectors produced by CyclicCAE to all-atom molecular structures, written to disk as Rosetta a silent file.

Optionally, `-out:file:silent _all_ atom structures.silent` may be replaced with `-out:file:o <prefix>` to directly write out PDB files with a given prefix.

#### S2.4 A Rosetta application for placing CyclicCAE-produced scaffolds for binder design: align _ml _backbones_ to _original_ target

In certain CyclicCAE design pipelines, it can be convenient to generate mainchain conformations with CyclicCAE (either using it for *de novo* conformational sampling or for de-noising some set of candidate conformations produced by some other means), convert them to PDB format or to Rosetta “silent” format with the `dof _vectors_ to _poses` application, and then to align them to stub residues docked to a target protein to design a binder to that protein. We wrote the `align _ml_ backbones_ to _original target` application to facilitate the alignment step. This application takes as input a target structure, a set of generated macrocycles, and a list of alignment residues, and produces as output a set of aligned target-macrocycle structures.

To use `align_ ml _backbones_ to _original_ target`, a user must first patch and compile Rosetta as described in section S2.1. A sample commandline command to run the application is shown in Supplementary Listing S14.

**Supplementary Listing S14.**
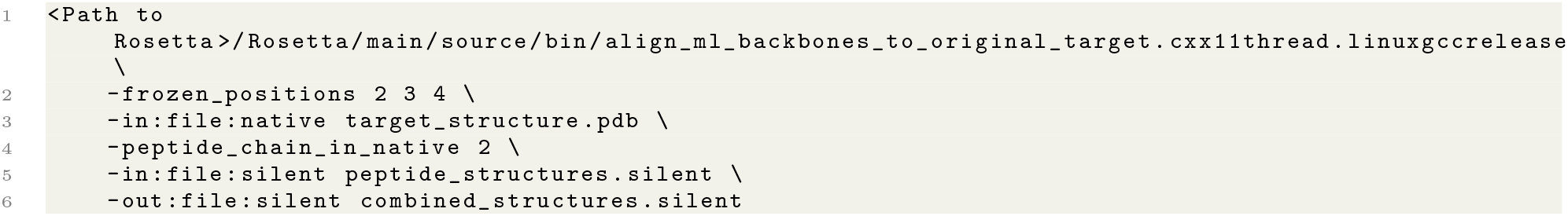
Commandline inputs to align a set of macrocycle scaffolds generated with CyclicCAE to a target protein structure.

In Supplementary Listing S14, the `-frozen positions` option is required, and lists the positions (based on *peptide* indexing) that are to be aligned with a docked peptide in the target structure. The target structure itself is provided with the `-in:file:native` option. The chain in this target structure that will be used for alignment and replaced with peptides is identified with `-peptide_ chain_ in_ native`. A value of 0 (the default value) means to use the last chain. Sampled peptide macrocycle conformations are provided as a Rosetta silent file with `-in:file:silent`, or alternatively as a set of PDB files with `-in:file:s` (with separate PDB files provided on the commandline) or with `in:file:l` (with a text file containing a list of PDB files provided on the commandline). The output structures will be written to the file specified with `-out:file:silent`.

### S3 CyclicCAE architecture and loss functions

#### S3.1 Conformational autoencoder neural network architecture

The conformational space accessible to poly-glycine peptide macrocycles spans all conformations accessible to chiral peptides made from any mixture of d- or l-*α*-amino acids. For small poly-glycine macrocycles (up to roughly 12 residues), this space may be more or less exhaustively sampled using `GeneralizedKIC`. Based on `GeneralizedKIC`‘s discrete sampling, our goal is to learn a continuous representation of the accessible low-energy peptide conformations. To this end, we introduce a mapping, *η*(·) : ℝ^*ND*^ → ℝ^*L*^, to convert the conformational degrees of freedom (DoFs) for an *N*-residue macrocycle with *D* degrees of freedom per residue into a much reduced *L*-dimensional latent space. Here, *η*(·) serves as an encoder function, while the inverse *η*^(−1)^ (·) : ℝ^*L*^ → ℝ^*ND*+*Q*^ serves as a decoder function that maps the latent space back to the input dimension, with *Q* additional outputs, as described below.

We sought to train an autoencoder neural net [20] with encoder and decoder limbs that could map an input degree-of-freedom vector 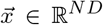 to a latent-space vector 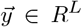, and then faithfully reconstruct a vector 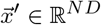:

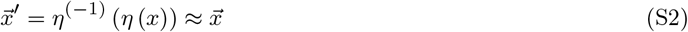

The architecture for this autoencoder neural network is shown in **Fig. 1** in the main text, with details shown in **Ext. Fig. 1**. Since torsion angles and bond angles are components of the degree-of-freedom vector 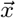, they presented a unique challenge compared to linear terms. Due to the circular nature of angles, where -180^°^ and 180^°^ are equivalent, we could not apply simple regression as with linear terms. To address this, we converted angles into radians and computed the sine and cosine of each one. This approach allowed regression or mean squared error (MSE) to be applied to the sine and cosine values, from which the original angle could be recomputed using the atan2(·, ·) function.

We used linear convolutional layers to capture correlations and patterns between degrees of freedom nearby in molecular connectivity, whether or not the atoms built by those degrees of freedom were close in space [21, 22, 23]. Dense multilayer perceptron (MLP) layers were used to compress relevant information into a reduced representation. Residual connections allowed important information from earlier steps to flow into the reduced representation, improving gradient flow and reducing the computational cost of training [24].

One unique feature of our architecture is that the decoder arm, *η*^−1^(·), maps *R*^*L*^ → *R*^*ND*+*Q*^, where *Q* is the dimensionality of a vector of energies, 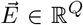. This 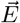 vector is predicted by the decoder using a series of MLP layers, in a parallel path alongside the MLP and deconvolutional layers that produce the decoded 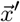 degree-of-freedom vector. The energies of the 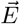 vector are *not* present in the original inputs. During training, the loss function penalizes both deviation of conformational degrees of freedom from the input conformation, as well as deviation of predicted energy values from known values (see section S3.2). During inference, the network’s ability to predict energy values can be used to compute gradients of energy with respect to latent space coordinates, and to thereby relax structures by gradient-marching in the latent space, remaining in the space of accessible conformations.

To permit efficient training, we wrote a Rosetta application, `extract_ dof _vectors`, which converts a set of mainchain conformations sampled with `GeneralizedKIC` to a binary database of records for input into a neural net. This is further described in section S2.2. Each record included the *ϕ, ψ*, and *ω* mainchain torsion angles, mainchain bond angles, and mainchain bond lengths, plus the energy of each structure as scored using Rosetta’s `ref2015` force field [19], the DFTB semi-empirical method [25], and the DFTB method with the `fa _sol`, `fa_ intra _sol _xover4`, and `lk_ ball_ wtd` implicit solvation terms from Rosetta’s `ref2015` added. The latter two energies made use of the `RosettaQM` extension (manuscript in preparation) which allows Rosetta to communicate with the `GAMESS` quantum chemistry package [13]. Notably, the training dataset did not include interatomic distances (or proxies like contact maps) for non-bonded atoms.

Since the terminal bond was not a DoF of the system, corrective terms were added to ensure ideal terminal bond lengths, terminal bond angles, and terminal dihedral angles, as shown in **Ext. Fig. 2** and described further in section S3.2.2.

#### S3.2 The loss function used in training

During training, we used a loss function composed of three primary components: the reconstruction loss 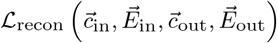, the terminal bond geometry loss 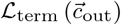, and the latent-space layout loss 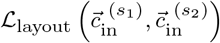. Given *N*_sam_ training samples in total or in a stochastic gradient descent batch, the overall loss function is given by Eq. (S3):

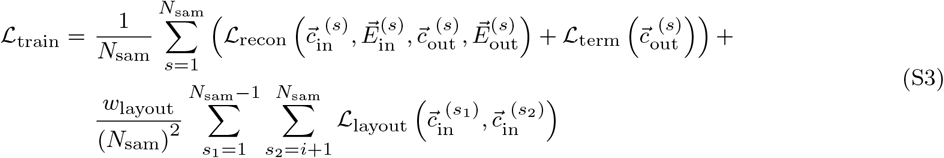

In Eq. (S3), 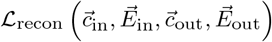 scores the reconstruction of an input conformation, expressed as an input DoF *N*_DoF_-vector 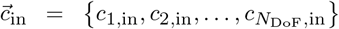, to produce an ouput DoF *N*_DoF_-vector 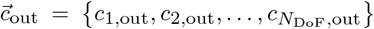. It also scores the ability to reproduce an input *N*_energy_-vector of energies, 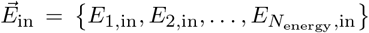, to produce an output *N* -vector of predicted energies, 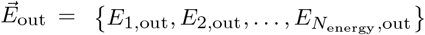. The terms contributing to the reconstruction loss are described in section S3.2.1.

The bond length, bond angles, and dihedral angle of the terminal amide bond are not part of the DoF vector defining the conformation, since these are fully determined by the bond lengths, bond angles, and dihedral angles of the rest of the macrocycle mainchain. Nevertheless, small errors in the mainchain DoFs can result in large errors in the terminal bond geometry due to lever-arm effects. For this reason, we included an additional term, 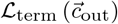, penalizing distorted geometry in the terminal amide bond on reconstruction. This is described in S3.2.2.

The final term, 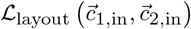, controls the latent-space arrangement, ensuring that similar conformations are nearby in the latent space while very different conformations are far apart. This is described in detail in section S3.2.3.

The relative weighting of these terms is described in section S3.5.

##### S3.2.1 The reconstruction loss terms

As shown in Eq. (S4), the reconstruction term of the losss function is broken down into terms representing reconstruction of bond lengths in the DoF vector, bond angles and dihedral angles in the DoF vector, and estimated energy.

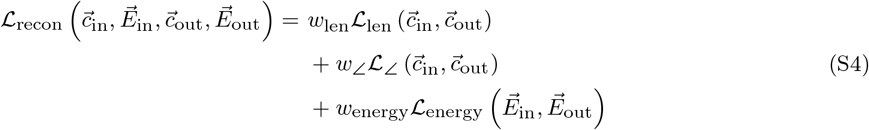

To describe each of these terms, we first define functions analogous to the Kronecker delta function. These terms are 1 if a given index in the DoF vector represents a DoF of a given type, and 0 otherwise, as shown in Eqs. (S5) and (S6):

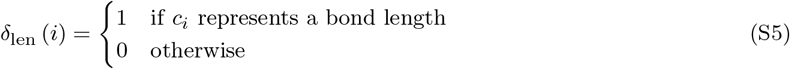

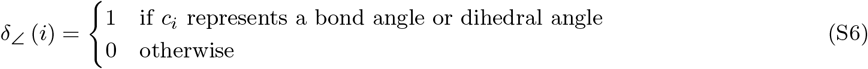

The bond length reconstruction term, a simple harmonic function of the difference between input and output bond lengths, is then given by Eq. (S7), where *N*_len_ represents the number of entries in the DoF vector representing bond lengths:

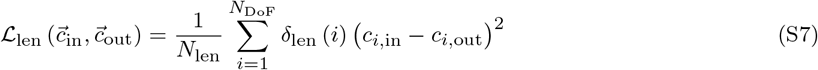

The bond angle and dihedral angle reconstruction term in turn is given by Eq. (S8), where *N*_∠_ represents the number of entries in the DoF vectors representing bond angles or dihedral angles:

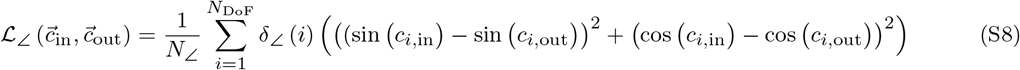

The energy reconstruction term is given by Eq. (S9). Like the bond length loss term, it is a simple harmonic penalty based on the difference between actual (input) energies and predicted (output) energies.

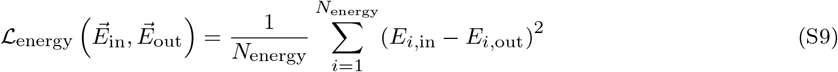

Each of these terms is given a weighting factor (*w*_len_, *w*_∠_, and *w*_energy_ in Eq. (S4)). These weights are described in section S3.5.

##### S3.2.2 The terminal amide bond geometry loss term

To penalize deviation from ideal terminal amide bond geometry, it was first necessary to compute the Cartesian coordinates of the atoms defining the terminal bond from the output DoF vector 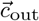. This required computing Cartesian coordinates for *each* atom within the poly-glycine macrocycle from DoF vector 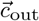. In a pre-computation, 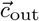 was divided into vectors 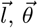, and 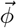, representing bond lengths, bond angles, and dihedral angles, respectively. The procedure for converting these to Cartesian coordinates of each mainchain atom is shown in Supplementary Algorithm S1.

###### Supplementary Algorithm S1

Procedure for determining coordinates of all macrocycle mainchain atoms from DoF vectors 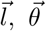, and 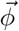, representing bond lengths, bond angles, and dihedral angles, respectively. In this, function 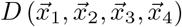, computes the dihedral angle between four points.

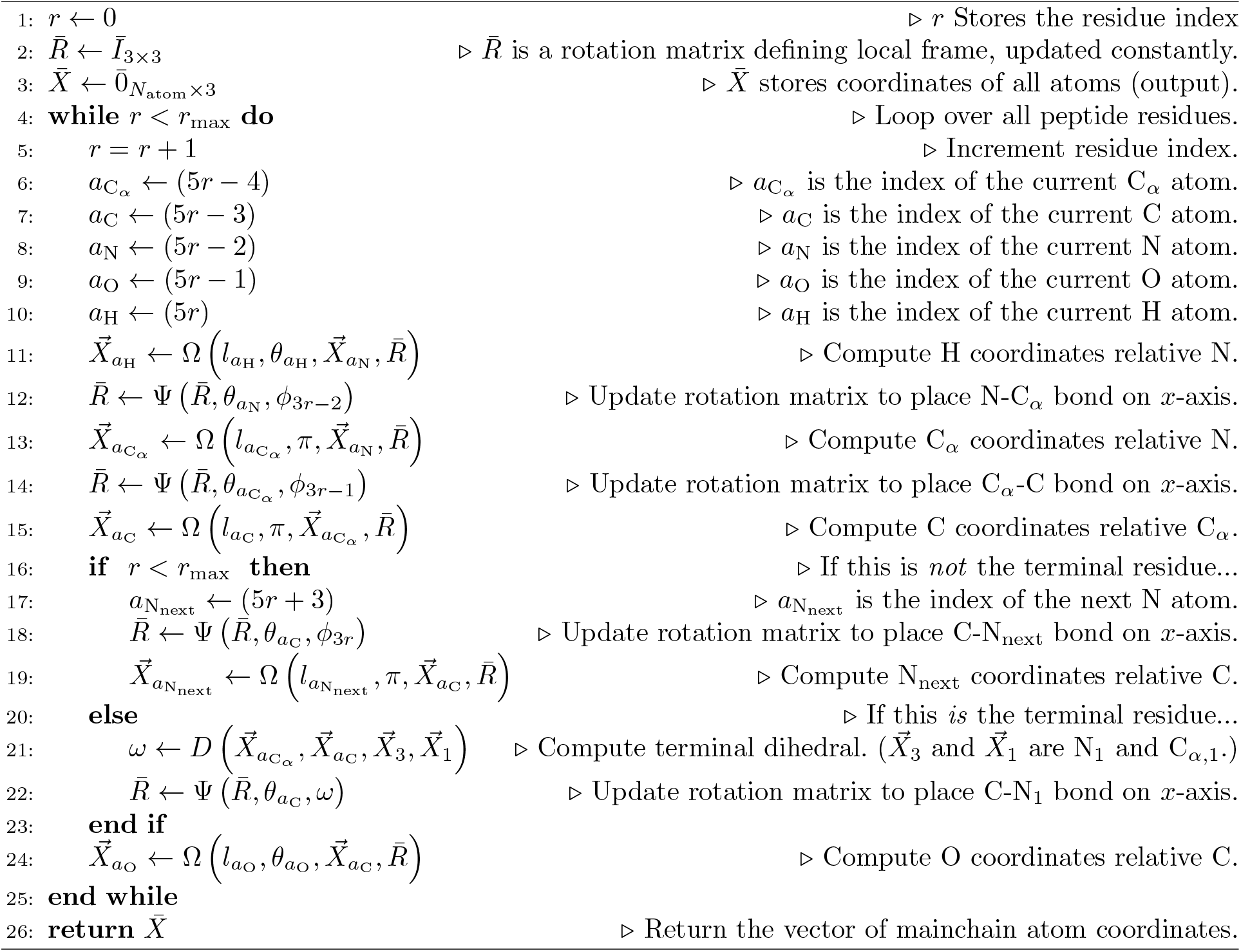

To calculate the Cartesian coordinates of each atom, we applied 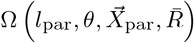, where *l*_par_ was the bond length to the atom’s parent, *θ* was the bond angle between the atom, its parent, and its grandparent in the parent’s reference frame (or *π* if the reference frame has been updated), 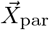 was the coordinate of the atom’s parent, and 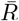 was the rotation matrix defining the reference frame of either the parent (for O or H atoms) or the current atom (for N, C_*α*_, or C atoms). This function is defined in Eq. (S10).

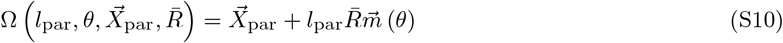

In Eq. (S10), the vector 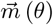 is given by Eq. (S11):

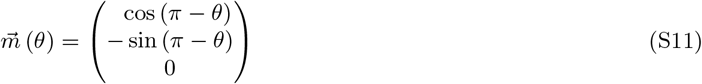

As shown in Supplementary Algorithm S1, the rotation matrix 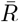 was initialized to the identity matrix. It was periodically updated by function 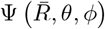, defined in Eq. (S12):

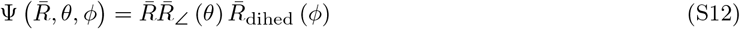

In Eq. (S12), the dihedral rotation matrix 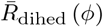 and the bond angle rotation matrix 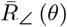 are given by Eqs. (S13) and (S14), respectively:

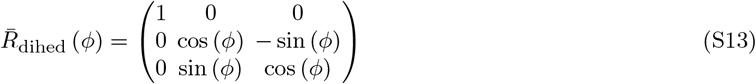

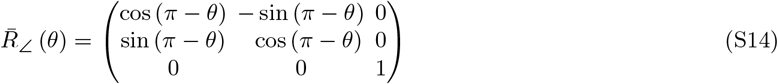

We represent Algorithm S1 as a function 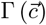 that converts a DoF vector 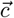 into mainchain atom coordinate matrix 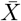.

The terminal loss penalty, 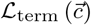, is given by Eq. (S15):

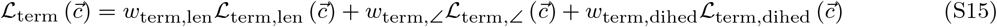

The first term in Eq. (S15) penalized deviation of the terminal bond length from an ideal value 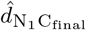 using a harmonic function, as shown in Eq. (S16):

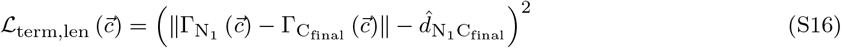

In Eq. (S16), we use 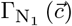 and 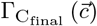 to represent the coordinates of the first residue’s nitrogen and the final residue’s carbon, as reconstructed from the DoF vector 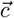, respectively. During training, ideal bond lengths 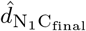 were taken from input structures; during inference, an ideal bond length of 1.36 Å was used.

The second term in Eq. (S15) penalizes deviation of the terminal bond angles ∠C_*α,i*_CN, ∠OCN, ∠CNH, and ∠CNC_*α,i*+1_ from their ideal values 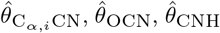, and 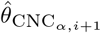, respectively. This is shown in Eq. (S17).

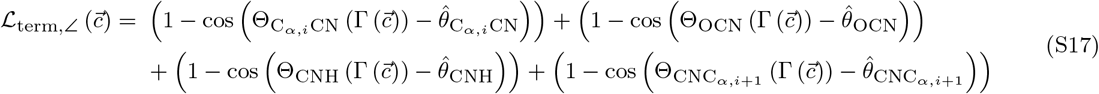

In Eq. (S17), the various 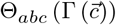 functions compute the bond angle ∠*abc* given the matrix of atomic coordinates produced from the DoF vector 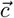 by function 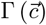. Ideal values of 116^°^, 123.0323^°^, 119.1399^°^, and 120^°^ were used for 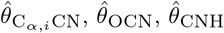, and 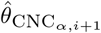, respectively during inference. The angle penalty term was not used during training (*i*.*e*. its weight *w*_term,∠_ was set to zero while training).

The final term in Eq. (S15) penalizes deviation from a *trans* amide bond conformation (a dihedral value differing significantly from 180^°^). It is given by Eq. (S18):

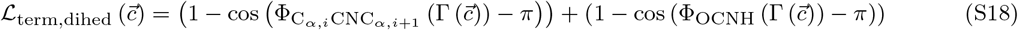

In Eq. (S18), 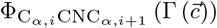 and 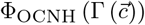 are functions that compute the C_*α,i*_CNC_*α,i*+1_ and OCNH dihedral angles, respectively, given the coordinate matrix produced by 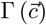 from the DoF vector 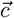. To reduce the expense of the calculation slightly, the trigonometric identitiy cos (*ϕ* − *π*) = − cos (*ϕ*) was employed in our implementation. Note that this term was also not used (*i*.*e*. the value of *w*_term,dihed_ was set to zero) during training.

##### S3.2.3 The latent space layout loss term

For the applications of CyclicCAE that we envisaged, a key feature needed to be that similar conformations map to nearby points in the latent space, and dissimilar conformations map to points distant from one another. This allows easy exploration of variants of an input structure that preserve favourable structural features such as long-range hydrogen bonding patterns, for instance by Monte Carlo methods (see section S4.6) or gradient descent minimization of predicted energy (see section S4.4). Although variational autoencoders and contractive autoencoders accomplish this, they tend to do so in a manner that imposes additional mathetmatical properties on the model which, while useful in certain circumstances, were not considerations in our case. We therefore chose to expand the loss function used in training with the latent-space “layout” term shown in Eq. (S19).

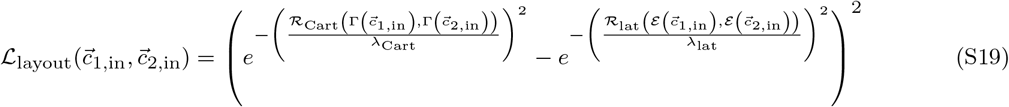

In Eq. (S19), 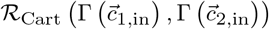 is the Cartesian-space mainchain heavy-atom root mean squared deviation (RMSD) function calculating the difference between the superimposed structures produced by DoF vectors 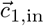 and 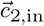 when reconstructed by the Γ (·) function, and 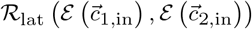 is the latent-space RMSD function computing the distance between points in the latent space. The encoder function mapping a DoF vector 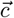 to a point in the latent space is represented by 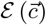. The user-defined parameters *λ*_Cart_ and *λ*_lat_ define the breadth of Gaussian functions in Cartesian and latent space respectively. These are used to define a concept of “closeness” in the two spaces: if two structures have a Cartesian-space RMSD that is small compared to *λ*_Cart_, then 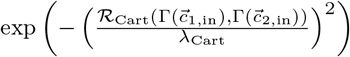 will evaluate to a value close to 1, while if it is large, it will evaluate to a value close to 0. Similarly, if two structures encode to points in the latent space that are separated by a distance that is small compared to *λ*_Cart_, then 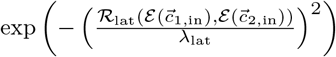 will evaluate to a value close to 1, while if this distance is large, it will be close to 0. The outer harmonic function constrains the two values to be similar. This ensures that gradient descent performed during training yields a model that maps similar structures (those with low Cartesian-space RMSD values) to nearby points in the latent space, and dissimilar structures (those with high Cartesian-space RMSD values) to distant points in the latent space.

As shown earlier in Eq. (S3), the full latent-space loss function involves iterating over all pairs of structures in the training dataset or in a stochastic gradient descent batch, making this function one of the more expensive functions to compute, since it scales quadratically with batch size. (This also necessitates the 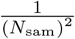 normalization in Eq. (S3)). To accelerate this calculation, the matrix of Cartesian-space RMSD values (the values produced by 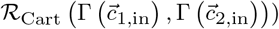) was pre-computed by the `extract _dof_ vectors` application (section S2.2).

#### S3.3 The loss function used during inference

Simple inference, as shown in **Fig. 1B** or **C**, involves no loss function. However, structures produced by CyclicCAE may be refined by gradient descent minimization (section S4.4) or Monte Carlo methods (section S4.6, and these do require a loss function. The loss function used for these inference tasks, 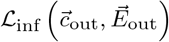, is given by Eq. (S20):

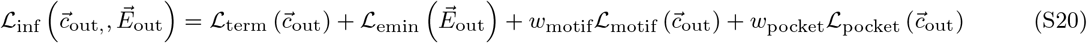

The first term in Eq. (S20), 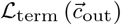, is a terminal geometry constraint term that is also used in the training loss function (Eq. (S3)). This was described in detail in section S3.2.2, above. Note that while nonzero weights were set only for the bond length component during training, during inference the weights on angle and dihedral constraints for the terminal bond are also nonzero. These are shown in Supplementary Table S5.

The second term is based on predicted energies, and is described in section S3.3.1. This permits geometry optimization to minimize predicted energy while preserving other favourable geometric features such as long-range hydrogen bonds.

The third term is used only when seeking scaffolds capable of presenting a known motif. This penalizes geometry unable to present this motif, and is described in section S3.3.2 (with further details about inpainting in section S4.7). When no motif is being used, the weight *w*_motif_ is set to zero.

Like the third term, the final loss term is specialized, used only when seeking scaffolds with shape complementarity to a target protein’s binding pocket. This term is modelled after the repulsive part of Rosetta’s Lennard-Jones potential [19], and is described in detail in section S3.3.3. When there is no binding pocket context, *w*_pocket_ is set to zero.

##### S3.3.1 The energy minimization loss term

Optimization of predicted energy is possible by including some weighted sum of the components of the predicted energy vector in the loss function, as shown in Eq. (S21).

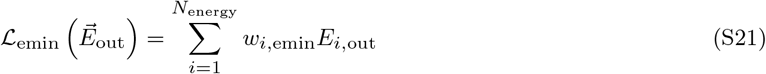

##### S3.3.2 The conformational motif loss term

If one wishes to carry out conformational motif inpainting with CyclicCAE, then one is seeking output torsion vectors in which a subset of the DoFs match the known DoFs of the linear motif. We define the DoF vector 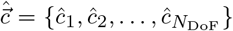, with entries whose DoF identities match the DoF vector 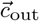 produced by the decorder limb of the CyclicCAE neural net. This vector’s entries match the DoFs of the motif where these align to the peptide, and are zero otherwise. In analogy to Eqs. (S5) and (S6), we define the delta function *δ*_motif_ (*i*), given by Eq. (S22):

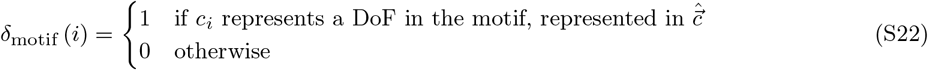

To find scaffolds with these DoFs, we introduce the loss function 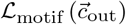, given by Eq. (S23):

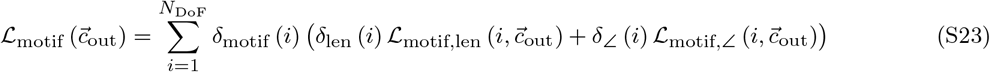

In Eq. (S23), 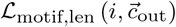 is a simple harmonic function constraining bond lengths in the peptide motif region to match those of the original motif, as shown in Eq. (S24):

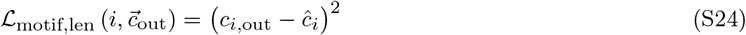

Similarly, 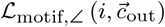 constrains bond angles and torsion angles in the peptide motif region to match those in the original motif, as shown in Eq. (S25):

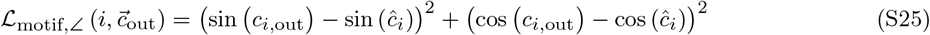

##### S3.3.3 The target binding pocket context loss term

CyclicCAE is trained on peptide backbones generated in isolation. In order to use CyclicCAE to relax peptide structures in the context of a binding pocket on a target protein, it is necessary to build in at least some of the physics of interaction between peptide and target. To this end, we added an additional term to the loss function used during Monte Carlo searches (section S4.6) and gradient-descent relaxation (section S4.4), 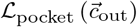, which represents the steric repulsion between target atoms and peptide atoms. To construct this loss function, we begin by taking the repulsive portion of the Lennard-Jones term from Rosetta’s *ref2015* energy function [19] to construct function *E*_rep_ *i, j, ε*_*i,j*_, *σ*_*i,j*_, representing the steric repulsion between a peptide atom with peptide atom index *i* and a target atom with target atom index *j*, as shown in Eq. (S26):

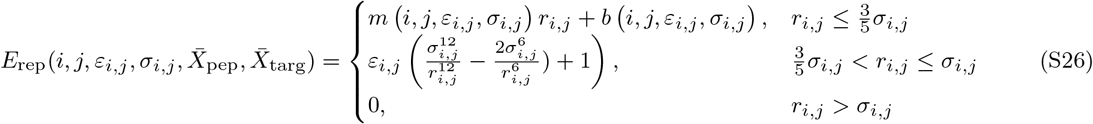

In Eq. (S26), 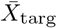 is an *N*__atom,targ__ *×*3 matrix of target heavy-atoms that should repel the peptide. CyclicCAE by default only considers target mainchain heavy-atoms plus C_*β*_ atoms, since subsequent design steps are likely to repack target side-chains and allow motions of atoms beyond the C_*β*_ atom. However, a user may optionally specify that all target heavy-atoms be included in the 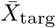 matrix. The value *r*_*i,j*_ is the Cartesian distance between atoms *i* in the peptide and *j* in the target, as shown in Eq. (S27).

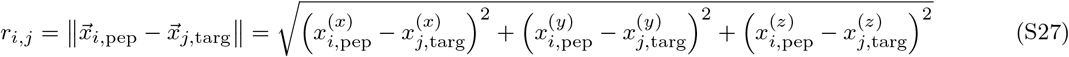

In Eq. (S26), *ε*_*i,j*_ and *σ*_*i,j*_ are parameters based on the types of atoms *i* and *j* which give the potential well depth and the combined Van der Waals radius, respectively. Values for these were taken from the `ref2015` force field [19]. The vectors 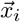 and 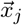 are the Cartesian coordinates of peptide atom *i* and target atom *j*, respectively. The values of *m i, j, ε*_*i,j*_, *σ*_*i,j*_ and *b i, j, ε*_*i,j*_, *σ*_*i,j*_ were chosen to ensure *C*^1^ continuity, and are given by Eqs. (S28) and (S29):

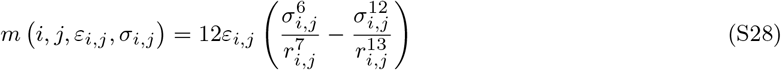

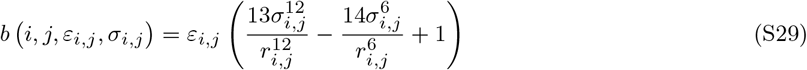

Since Eq. (S26) depends on Cartesian coordinates of peptide atoms, it is necessary to generate these from the DoF vector 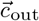 produced by CyclicCAE, and to align these to the target using coordinates of user-designated “stub” residues already placed in the binding pocket. The former is accomplished by applying function 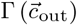, described in section S3.2.2. The latter is achieved using the Kabsch-Umeyama algorithm [26].

To apply the Kabsch-Umeyama algorithm, we must first mask the peptide atoms that are not involved in the alignment. In analogy with Eqs. (S5), (S6), and (S22), we define function *δ*_stub_ (*i*) to indicate whether a given peptide atom is one involved in alignment or not:

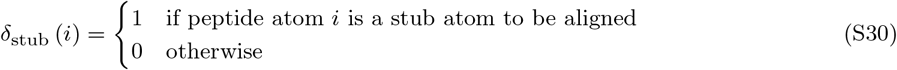

Given CyclicCAE’s Cartesian output coordinates 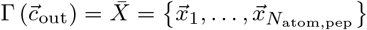, and the corresponding user-provided stub coordinates 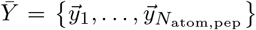 (where the various 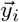 vectors are nonzero only for the subset of atoms in the stub), we begin by computing the geometric centroids of the stub atoms in both sets:

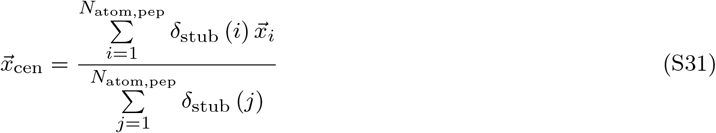

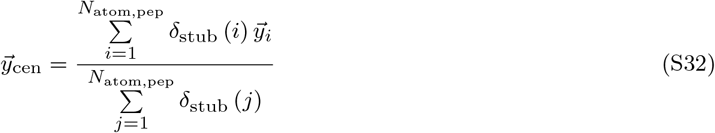

Then, we centre all peptide stub atom coordinates and user-provided stub atom points around their respective centroids. We now define *N*_atom,stub_ as the number of stub atoms, and the mapping *ν* : ℕ → ℕ to map peptide indices to stub indices (for peptide atom indices that represent stub atoms), and we prepare the matrices 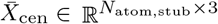 and 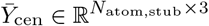:

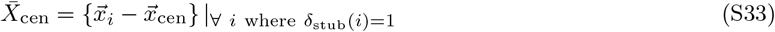

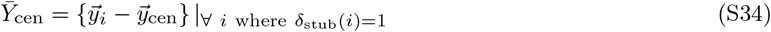

Next, we construct the covariance matrix 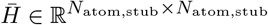, as shown in Eq. (S35):

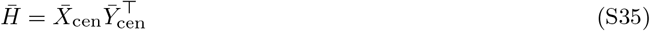

We then apply singular value decomposition (SVD) to covariance matrix 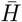 to obtain *Ū* ∈ ℝ^3*×*3^, singular value vector 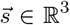, and conjugate transpose rotation matrix 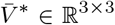 which minimizes the RMSD between 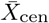 and 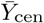, as shown in Eq. (S36):

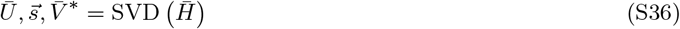

To overlay the peptide points onto the stub atoms, the rotation matrix, 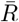, may now be calculated from the *Ū*and 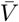 matrices, as shown in Eq. (S37), while the translation vector 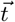 can be computed from the difference in the centroids, as shown in Eq. (S38):

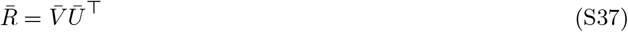

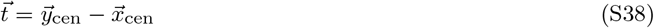

If 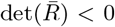, we flip the sign of the last column of 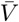, then reapply Eq. (S37) to recalculate 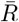. Finally, we align the peptide points 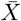 by applying the rotation 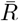 and translation 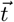 to produce the aligned points 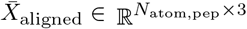. We use 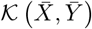 to represent the Kabsch-Umeyama algorithm giving the alignment of peptide atoms, 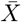, to a subset of stub atoms with coordinates given in 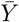.

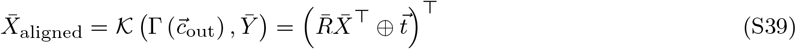

In Eq. (S39), the ⊕ symbol is used to represent element-wise column vector addition (addition of the vector to each column of the matrix). Given the aligned coordinates of peptide atoms, the coordinates and identities of target atoms, and the well depths (*ε*) and Van der Waal pair distances (*σ*) from the Rosetta `ref2015` energy function, it is possible to evaluate 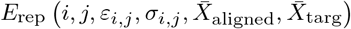, and, by extension, the loss function 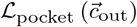, shown in Eq. (S40):

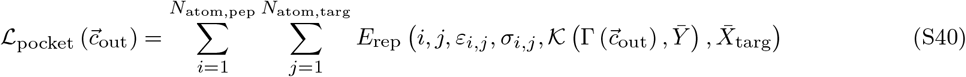

**Supplementary Table S5.** Weights of loss terms used during training and during gradient descent refinement or Markov chain Monte Carlo searches during inference. A dagger (*†*) indicates weights that are nonzero only when a particular feature (energy minimization, motif reproduction, or pocket-context repulsion) is used. “N/A” indicates that a term is not applicable in a training or inference context.

| Loss term | Weight parameter | Training value | Default inference value | Equation |
| --- | --- | --- | --- | --- |
| Latent space layout | $w_{\text{layout}}$ | 100.0 | 0.0 (N/A) | Eq. (S3) |
| Bond length reconstruction | $w_{\text{len}}$ | $1.0 \times 10^3$ | 0.0 (N/A) | Eq. (S4) |
| Bond/dihedral angle reconstruction | $w_{\angle}$ | $1.0 \times 10^5$ | 0.0 (N/A) | Eq. (S4) |
| Energy prediction accuracy | $w_{\text{energy}}$ | $1.0 \times 10^4$ | 0.0 (N/A) | Eq. (S4) |
| Terminal bond length | $w_{\text{term, len}}$ | 4.0 | 4.0 | Eq. (S15) |
| Terminal bond angles | $w_{\text{term, } \angle}$ | 0.0 | 1.0 | Eq. (S15) |
| Terminal bond dihedral | $w_{\text{term, dihedral}}$ | 0.0 | 0.5 | Eq. (S15) |
| Motif reproduction | $w_{\text{motif}}$ | 0.0 (N/A) | 1.0<sup>†</sup> | Eq. (S20) |
| Pocket Lennard-Jones repulsion | $w_{\text{pocket}}$ | 0.0 (N/A) | 1.0<sup>†</sup> | Eq. (S20) |
| Energy minimization | $w_{i, \text{emin}}$ | 0.0 (N/A) | 1.0<sup>†</sup> | Eq. (S21) |

#### S3.4 Loss function differentiability for training and latent-space gradient-descent relaxation

The loss functions used in training and in inference are implemented in the Pytorch framework [27], and are fully end-to-end differentiable using Pytorch’s automatic differentiation tools. This allows the calculation of the training loss gradient with respect to model parameters, to allow training. This also permits computation of gradients of the inference loss function with respect to latent-space coordinates, allowing gradient marching within the latent space. Since this allows macrocycle backbones to be relaxed in a less rugged, lower-dimensional space, it permits larger relaxations that still preserve important structural features such as internal hydrogen bond networks. When the target binding pocket context term is included (section S3.3.3), it also allows relaxation to relieve clashes with the target.

#### S3.5 Weights of loss terms used during training and inference

Weights of loss terms were tuned approximately by trial-and-error, seeking values that yielded trained models without egregious bias in the types of errors seen. More rigorous balancing is likely possible. Final weights used for training and for inference are shown in Supplementary Table S5.

### S4 Obtaining, training, and using CyclicCAE

#### S4.1 Obtaining the CyclicCAE architectural code and training scripts

CyclicCAE is publicly released under a free and open-source AGPL 3.0 licence. It may be obtained by cloning the CyclicCAE repository from GitHub (https://github.com/flatironinstitute/CyclicCAE_public). Commands for doing this are shown in Supplementary Listing S15.

**Supplementary Listing S15.**
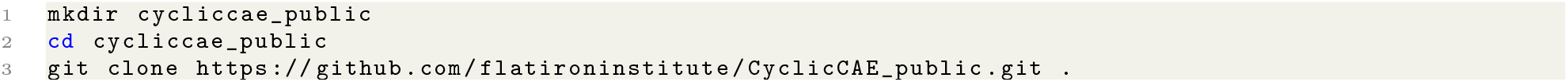
Command-line commands for cloning the CyclicCAE Git repository. In the final line, note the full stop at the end, indicating the current directory.

#### S4.2 Obtaining weights for pre-trained CyclicCAE models

Weights for pre-trained CyclicCAE models trained on 6-, 7-, 8-, and 9-residue poly-glycine peptide macrocycles are provided in a Git LFS (*L*arge *F* ile *S* torage) repository, with pointers in the main CyclicCAE repository, located in the `weights/` sub-directory. If a user has Git LFS installed on their system, then the cloning command shown above in Supplementary Listing S15 should also pull in the weights. If Git LFS is not installed, or if the `weights/` directory contains only small pointer files after cloning the CyclicCAE directory, then a user should follow the instructions at https:gitlfs.com to install Git LFS. Once this is done, the command shown in Supplementary Listing S16 will retrieve the weights.

**Supplementary Listing S16.**
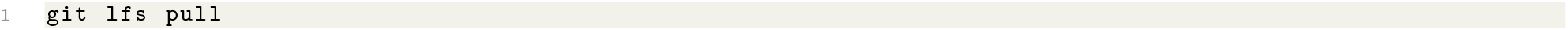
Command-line command for obtaining the CyclicCAE pre-trained model weights using Git LFS. Note that Git LFS must first be installed and activated.

#### S4.3 Building and training CyclicCAE models

In order to train CyclicCAE one needs to construct a torch `Tensor` object, 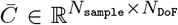, where N_sample_ is the number of conformations in the training set, and *N*_DoF_ is the number of degrees of freedom (DoFs) in each conformation. Required DoFs are torsion, bond angle, and bond lengths for each atom, which can be extracted using `Rosetta` or `PyRosetta` (see section S2.2). An example using `PyRosetta` is shown in Supplementary Listing S17:

**Supplementary Listing S17.**
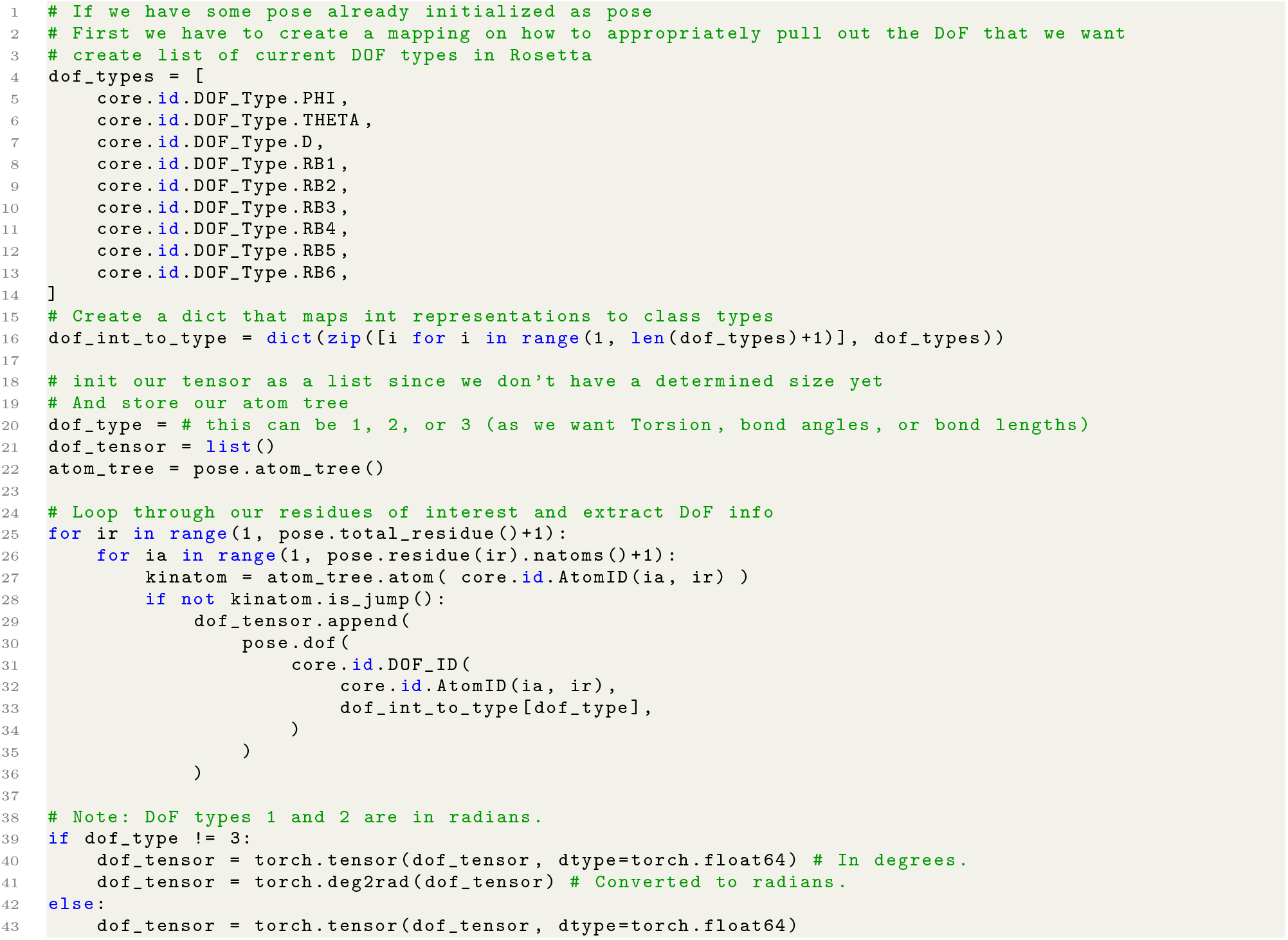
Example Python script for training CyclicCAE.

For each DoF extracted, it is then necessary to concatenate the different types of DoFs to construct a vector of length *N*_DoF_. It is also necessary to prepare a matrix of energies to allow the model to learn to predict energies. In our case, in addition to using Rosetta’s `ref2015` force field [19], we also used `RosettaQM` (manuscript in preparation) to compute eneriges using quantum chemistry calculations performed with `GAMESS` [13]. We used the DFTB semi-empirical method [28] with organic solvent (benzene) used with the SMD implicit solvation model [29]. We used this solvation model to avoid complications that arise from the hydrophobic effect when trying to model implicit aqueous solvent. Three energies were appended to the end of the *N*_DoF_-vector: the `ref2015` force field energy, the DFTB energy computed with `RosettaQM` and `GAMESS`, and the DFTB energy plus the solvation terms (`fa sol`, `fa_ intra_ sol_ xover4`, and `lk_ ball _wtd`) from the `ref2015` force field (an experiment in using QM calculations to capture the enthalpic component of conformational energy while using Rosetta’s force field for aqueous solvent entropy estimates).

The final needed input is a parameter map indicating which DoFs represent torsions, bond angles, bond lengths, or energies. This is added to the `params` map which is saved with the input dataset. An example for 6-mers can be found in Supplementary Listing S18:

**Supplementary Listing S18.**
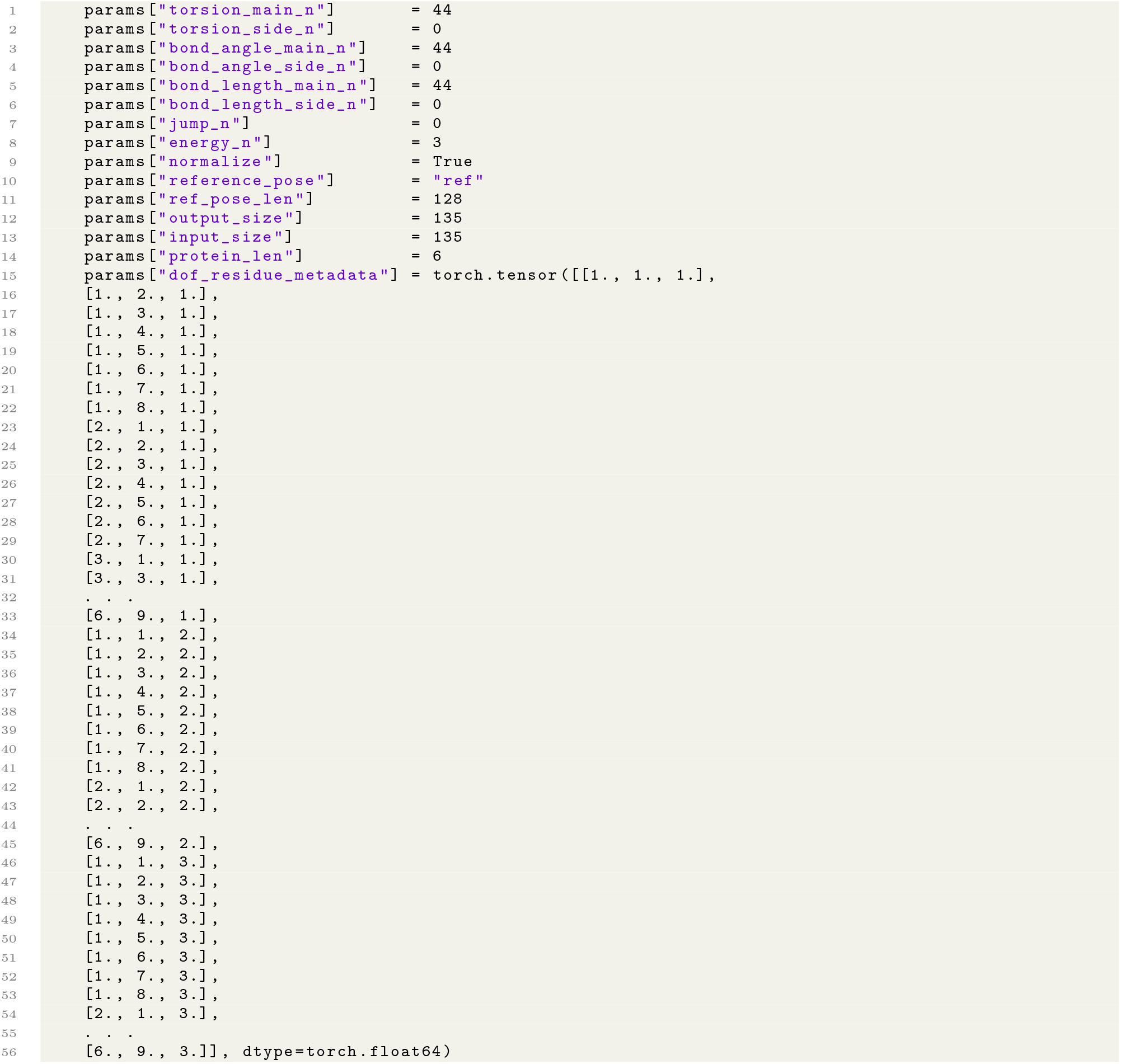
Example parameters map for training CyclicCAE on a 6-mer training dataset.

The total count of torsions, bond angles, and bond lengths is equal to the number of atoms present with the addition of 3 virtual atoms, added for closure. Here `jump_n` is zero, since we are only looking at a monomer in isolation. Both `input_ size` and `output_ size` represent the length of the combined DoF vector. In order to map the produced DoFs back to the residue DoFs accurately, we use a `dof _residue _metadata` tensor. This stores the residue index (first column), atom index (second column), and an integer *d* ∈ *{*1, 2, 3*}* representing the DoF type (third column), where 1 represents a dihedral, 2 represents a bond angle, and 3 respresents a bond length. Root atoms are omitted from the metadata tensor.

In addition to the matrix 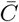, containing all training data, we also prepared a reduced training data matrix 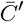. This was prepared by clustering sampled conformations using the `energy_based_clustering` application (see section S1.4) and using the ten lowest-energy conformations in each cluster.

With input matrices 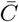 and 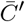, CyclicCAE can be trained. First we carried out an initial round of pretraining to get the model to approximate 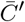, yielding 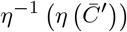, followed by refinement on the full input data

**Supplementary Table S6.**
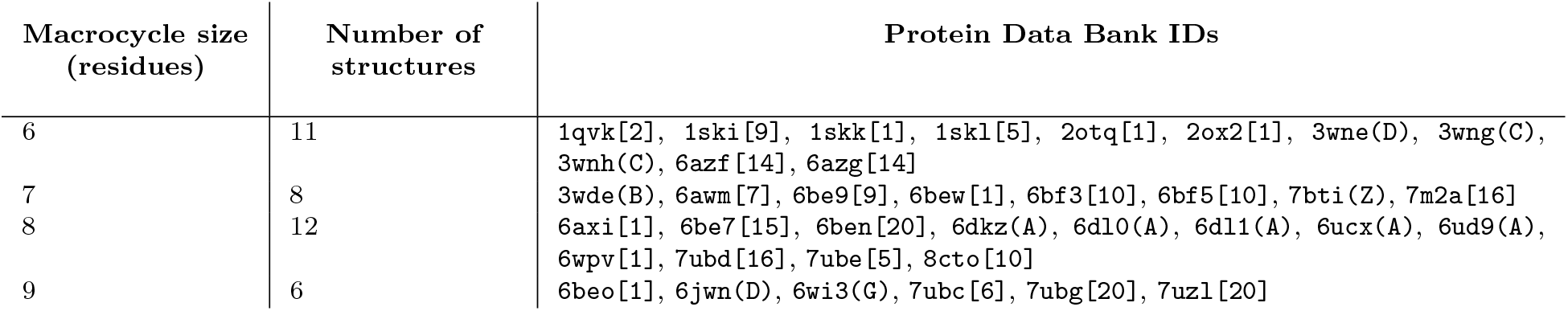
Peptide macrocycles from the PDB used to benchmark CyclicCAE’s ability to reproduce known structures. For cases of x-ray crystal structures in which multiple chains are present in the PDB file, the chain ID used is shown in round brackets. In the case of NMR structures, the structure in the ensemble that was used is shown in square brackets.

Set 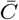, yielding 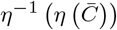. We trained the model using the *model*.*py* script with arguments of 3000 epochs, a latent-space size of 6, linear positional encodings, convolutional processing, a noise standard deviation of 1, noise iterations of 3, dropout of 0.1, and an initial learning rate of 1.0 *×* 10^−3^. Batch size was dependent on the size of *N*. For most of the pre-training, since *N* ≤ 3000, we used a batch size of 128 to get a somewhat similar number of scheduler steps per epoch per dataset. For the larger 9-mer set, we had to increase the batch size to 512 for the 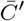 pre-training. The full training dataset was trained using the best weights, based on validation loss, with the same training schema as above. However, the batch size was increased to 2,064 for all models except for the 9-mers, which were trained with a batch size of 7,024 per GPU.

Please see the `README.md` file in the CyclicCAE Git repository for commandline options for training CyclicCAE.

#### S4.4 De-noising or relaxing input structures with CyclicCAE

A peptide in PDB format can be passed to CyclicCAE, which will extract the peptide’s mainchain degrees of freedom and pass them through the encoder and decoder, converting the peptide’s conformation to the closest conformation represented within the latent space. This conformation can then optionally be relaxed using the model with gradient descent, where the objective function being minimized is the conformational energy estimated by CyclicCAE. Since this objective function is an approximation of `ref2015`, this minimization approximates the minimization that Rosetta performs with its `MinMover` module. Supplementary Listing S19 shows an example commandline for de-noising with gradient descent relaxation:

**Supplementary Listing S19.**
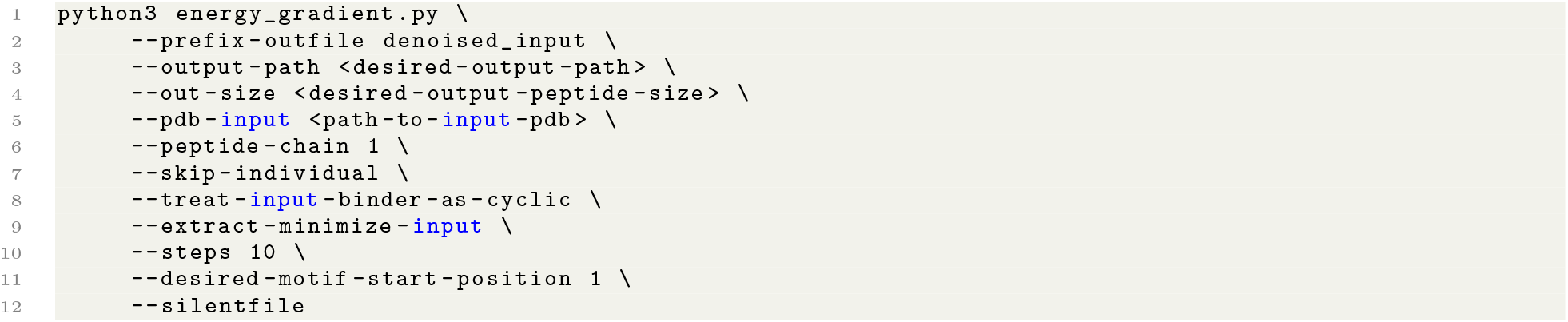
Commandline inputs to de-noise an input structure with CyclicCAE, with 10 steps of subsequent gradient descent relaxation based on estimated `ref2015` energy.

Please see the file `README.md` in the CyclicCAE Git repository (https://github.com/flatironinstitute/CyclicCAE_public) for full documentation on denoising input structures using the `--pdb-input` commandline flag, or on carrying out energy minimization using the `--extract-minimize-input` option.

We carried out de-noising on all available peptide macrocycle structures found in the PDB, subject to the restrictions described in the Methods section in the main text. The de-noised structures are listed in Supplementary Table S6.

#### S4.5 Sampling plausible “designable” macrocycle conformations with CyclicCAE

CyclicCAE’s latent space represents all plausible conformations of a peptide macrocycle of a given size, with plausibility based on the training dataset. Sampling points randomly from the latent space, and passing these through the decoding limb of the neural network, produces plausible conformational samples. Depending on the application, these may be used directly or further refined by gradient descent minimization in the latent space, using estimated energy as the objective function. Commandline inputs that carry out random sampling plus gradient descent minimization are shown in Supplementary Listing S20.

**Supplementary Listing S20.**
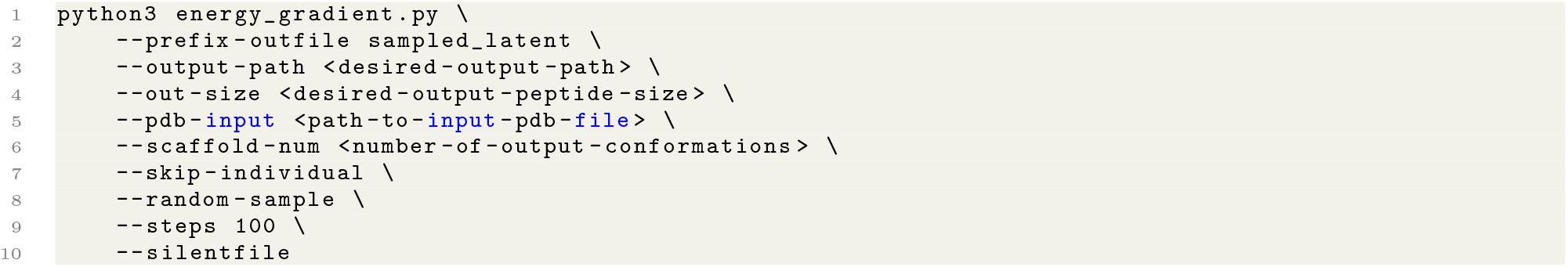
Commandline inputs to directly sample from the CyclicCAE latent space, with 100 steps of subsequent gradient descent relaxation based on estimated `ref2015` energy.

Please refer to the `README.md` file in the CyclicCAE Git repository (https://github.com/flatironinstitute/CyclicCAE_public) for full documentation on randomly sampling with the `--random-sample` commandline flag.

#### S4.6 Markov chain Monte Carlo latent-space exploration

CyclicCAE provides a bidirectional mapping from a high-dimensional conformational space to a low-dimensional latent space. When a particular peptide conformation is encoded as a point in the latent space, the local neighbourhood represents structurally similar peptide conformations thanks to the 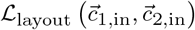 term included in the loss function during training (described in section S3.2.3). Markov chain Monte Carlo (MCMC) sampling can be used to explore this local latent-space neighbourhood in order to generate macrocycle conformations similar to a starting point. Alternatively, if a user desires more structurally diverse outputs, larger-scale MCMC trajectories can be used to explore more distant regions on the manifold, far from the original point. **Fig. 1D** in the main text illustrates this concept. Note that although MCMC can also be usd to construct thermodynamic distributions, here we are using it as a convenient way of finding low-energy or low-scoring conformations rapidly.

##### Supplementary Algorithm S2

Procedure for exploring latent space by MCMC to find a conformation 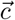 minimizing a loss function 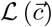.

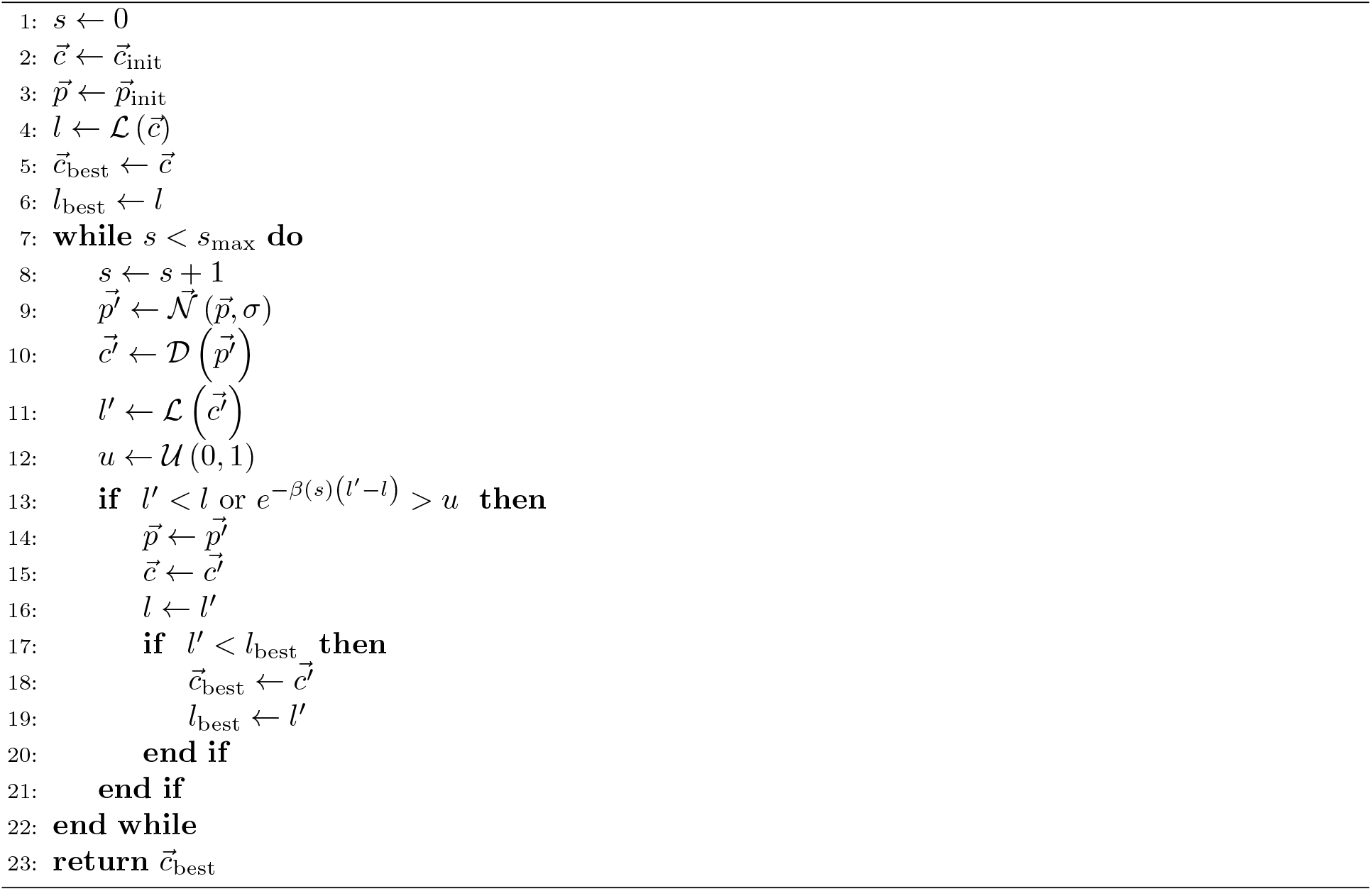

Given a starting conformation 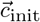 and the corresponding latent-space point 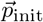, a loss function *L*, and the CyclicCAE decoder function *D*, the algorithm for MCMC-based latent-space sampling is shown in Supplementary Algorithm S2. In this, 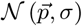 represents a random vector with the same dimensionality as 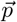 and with components *N*_*i*_ drawn from Gaussian distributions with centre *p*_*i*_ and breadth *σ*, and *U* (0, 1) is a random real number drawn uniformly in the interval [0, 1]. The thermodynamic beta, *β* (*s*), is a user input that governs the balance between hill-climbing for exploration and valley-descent for local optimization. This may be ramped over the trajectory if one wishes to perform simulated annealing.

With CyclicCAE’s `energy_gradient.py` module, MCMC-based latent-space exploration is possible using the `--mcmc` commandline flag. Full documentation for running Markov chain Monte Carlo latent-space searches with CyclicCAE is found in the CyclicCAE Git repository (https://github.com/flatironinstitute/CyclicCAE_public), in the `README.md` file.

#### S4.7 Using CyclicCAE for cyclization of linear sequences by conformational motif inpainting

Given a linear sequence taken from a known binding region of a protein, or a linear peptide that binds to a target protein, one may wish to design a peptide macrocycle that stabilizes this sequence in the binding-competent conformation. If the linear sequence’s target is of therapeutic interest, this can be a path to making a peptide macrocycle drug. This means finding plausible geometry for a few residues of peptide mainchain connecting the ends of the linear motif, so that the result is a plausible peptide fold that can be further stabilized by appropriate choice of the sequence of the added residues. While the second step may be performed with Rosetta’s `FastDesign` mover, the first step is well-suited to conformational motif inpainting using CyclicCAE.

To carry out conformational motif inpainting with CyclicCAE, we define the loss term 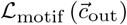, as shown in Eq. (S23) in section S3.3.2, and construct the DoF vector 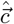 described in the same section by extracting DoFs from the motif. We then randomly sample the latent space for structures with similar DoFs to the selected residues. Starting with a predetermined number of scaffold structures, we use Markov chain Monte Carlo sampling (see section S4.6) with the motif loss term 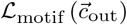 to identify mainchain conformations able to present the desired motif. Once an initial set of scaffolds is identified, we apply gradient-descent loss function minimization for *s* steps, using the full 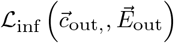 loss function shown in Eq. (S20) in section S3.3, which includes both energy and motif terms, to relax the structure while simultaneously maximizing the match to the motif of interest.

CyclicCAE’s `energy_gradient.py` module permits motif inpainting with the `--inpaint` commandline flag. Full documentation for running motif inpainting with CyclicCAE is found in the CyclicCAE Git repository (https://github.com/flatironinstitute/CyclicCAE_public), in the `README.md` file.

